# Non-preferred contrast responses in the *Drosophila* motion pathways reveal a receptive field structure that explains a common visual illusion

**DOI:** 10.1101/2021.04.29.442028

**Authors:** Eyal Gruntman, Pablo Reimers, Sandro Romani, Michael B. Reiser

## Abstract

Diverse sensory systems, from audition to thermosensation, feature a separation of inputs into ON (increments) and OFF (decrements) signals. In the *Drosophila* visual system, separate ON and OFF pathways compute the direction of motion, yet anatomical and functional studies have identified some crosstalk between these channels. We used this well-studied circuit to ask whether the motion computation depends on ON-OFF pathway crosstalk. Using whole-cell electrophysiology we recorded visual responses of T4 (ON) and T5 (OFF) cells and discovered that both cell types are also directionally selective in response to non-preferred contrast motion. We mapped T4s’ and T5s’ composite ON-OFF receptive fields and found they share a similar spatiotemporal structure. We fit a biophysical model to these receptive fields that accurately predicts directionally selective T4 and T5 responses to both ON and OFF moving stimuli. This model also provides a detailed mechanistic explanation for the directional-preference inversion in response to a prominent visual illusion, a result we corroborate with electrophysiological recordings and behavioral responses of flying flies.

## Introduction

In both invertebrate and vertebrate visual systems, neuronal signals bifurcate into parallel pathways that preferentially encode luminance increments (ON) or luminance decrements (OFF) within 1-2 synaptic layers of photoreceptors [1]. The direction of local motion is computed separately within these pathways a few synapses further downstream of the ON-OFF split [1–3]. Splitting sensory signals into increments and decrements is common to different modalities [4–6] and may enable more efficient stimulus encoding [7]. However, in the mammalian retina, motion is also computed in ON-OFF cells [8], and the separate motion pathways in the fly show clear evidence of crosstalk [9,10]. What is the benefit of mixing between pathways, and is it integral to computing motion? A potent tool for studying this question is the ‘reverse-phi’ visual illusion— perceived by both invertebrates and vertebrates [11–15]—in which inverting the contrast of moving objects (bright pixels become dark and vice versa) induces an illusory inversion in the detected motion direction [11]. Similarities between flies and vertebrates in bifurcating ON-OFF pathways as well as susceptibility to directional preference inversion when challenged with ON-OFF mixtures, suggest that understanding ON-OFF crosstalk in the *Drosophila* motion circuit could reveal fundamental aspects of visual processing, and may uncover more general, conserved aspects of sensory processing.

Anatomical studies of the *Drosophila* medulla identified two major pathways [16,17], that were later functionally determined as one preferentially encoding ON signals and another preferring OFF signals [18–20]. Each pathway contains motion computing circuits; the T4 neurons (ON pathway) and T5 neurons (OFF pathway) are the first cells in the visual system to show directionally selective responses [21]. Connectomic reconstructions mapped the T4 and T5 inputs and validated the proposal of separated pathways, with most connections between neurons of each pathway. However, anatomical interactions between prominent cells of the ON and OFF pathways have been described [9,17,22], whose contributions may be measured by recent functional studies [23]. For example, a key T4 input is primarily OFF-responding [24,25] and T5 neurons show responses to some ON stimuli [26]. The classic experiments that established the algorithmic analysis of motion computation in insects already showed perceptual inversions to ON-OFF combinations related to the reverse-phi illusion [13]. Later studies found neuronal correlates of this behavioral inversion in neurons downstream of T4 and T5 [12,27], and recent studies showed that T4 and T5 exhibit an inverted directional preference to reverse-phi stimuli [28]. Localizing this inversion to the directionally selective neurons suggests that mixing ON and OFF signals is a central and underappreciated feature of computing motion, but it is not known why or how T4 and T5 neurons are susceptible to the reverse-phi illusion.

In previous work, we described the mechanism by which T4 and T5 neurons generate directionally selective responses to moving bright and dark objects, respectively [29,30]. Using static stimuli of the Preferred Contrast (PC, bright for T4, dark for T5), we showed that both cell types share a similar spatiotemporal receptive field. This was unexpected since their input neurons have quite different properties [9,25,31]. We modelled this structure as generated by a fast excitatory conductance and a slower, spatially-offset inhibitory conductance [29,30]. In pilot experiments, we found that T4 and T5 cells generated directionally selective responses also to moving stimuli of their Non-preferred Contrast (NC, dark for T4, bright for T5). No contemporary understanding of how these cells generate directionally selective responses could account for these surprising observations. Therefore, in this study, we sought to understand how NC inputs affect T4 and T5 responses, and how these responses become directionally selective. First, we characterized T4 and T5 responses to PC and NC stimuli, and mapped their detailed receptive fields using both bright and dark stimuli. This revealed that T4 and T5 cells share a similar NC spatial receptive field. Next, we proposed a unified, conductance-based model to capture the PC/NC receptive field, and showed this model can explain the generation of directionally selective responses to both PC and NC moving stimuli. We found that this model also provides an explanatory mechanism for the reverse-phi illusion. Finally, we corroborate these model predictions with functional recordings and behavioral responses to reverse-phi motion and show that NC responses and their associated components are essential for understanding the motion computation.

## Results

### T4 and T5 generate directionally selective responses to non-preferred contrast stimuli

We used targeted in-vivo whole-cell electrophysiology to localize the receptive field center for individual T4 or T5 neurons, identify their primary motion axis, and record responses to a panel of receptive field referenced stimuli (Fig. 1A). Throughout, we refer to bright over intermediate intensity background as bright stimuli, and dark over intermediate background as dark stimuli (see Methods). Bright stimuli are the PC for T4 but NC for T5, while dark stimuli are T5’s PC and T4’s NC (Fig. 1B). When T4 cells were presented with bright bars moving in the preferred and non-preferred directions (PD and ND, respectively), their responses were directionally selective (Fig. 1C_i_, top), as expected from prior work [21,29,30,32]. When presented with fast moving (56°/sec) dark narrow bars, T4 were largely unresponsive, regardless of the movement direction (Fig. 1C_i_, bottom). However, when presented with 4x wider dark bars moving at the same speed, T4 responses were large and directionally selective (Fig 1. C_i_, bottom right). T5 cells showed similar responses to bright moving stimuli: narrow bars only evoked small responses, while wider bars evoked larger, directionally selective responses (Fig. 1C_ii_, bottom right). We further explored T4 and T5 responses to NC bars of different widths moving at different speeds, and found that responses to fast, narrow NC bars were not directionally selective, while responses to slow, wide NC bars were. The difference between the peak PD and ND responses (proxy for directional selectivity) in these conditions is comparable in magnitude to this difference for PC bars of the same size and speed (Fig. 1D).

**Figure 1:**
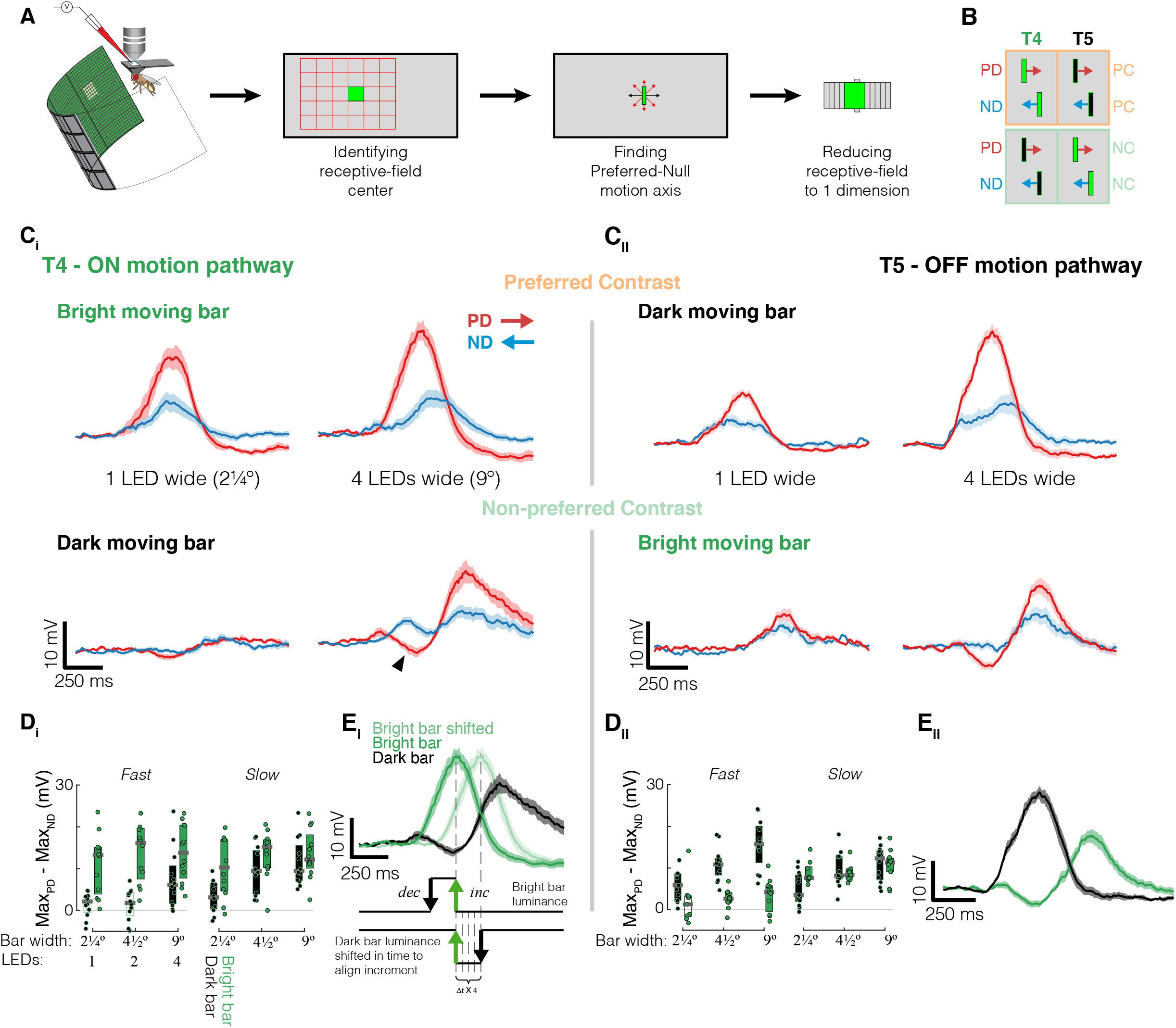
T4 and T5 generate directionally selective responses to non-preferred contrast stimuli. (A) Schematic of experimental setup and procedure. Whole cell recordings were targeted to GFP-labelled T4 and T5 somata (in flies with different genotypes, see Methods). The receptive field center for each cell was located by presenting a grid of flashing squares of the expected preferred contrast. The preferred direction was determined by presenting moving bars in 8 different directions. Further receptive field characterization was reduced to presenting stimuli along the PD-ND axis. (B) Summary of stimulus conventions used throughout. PD: Preferred Direction, ND: Nonpreferred Direction, PC: Preferred contrast, NC: Non-preferred Contrast. By convention, in all figures, the PD is aligned to movement from left to right regardless of the PD of each recorded cell. (C_i_) Mean baseline subtracted T4 responses (n=9 cells diagonally aligned PD-ND axis) to bars of width 1 and 4 LEDs (2.25° and 9°) moving at 56°/s (40ms per LED step) in the preferred and non-preferred direction with either preferred (top) or non-preferred (bottom) contrast. (C_ii_) Same as C_i_ for T5 (n=12 cells diagonally aligned PD-ND axis). (D_i_) Boxplots for difference in the peak response to motion in both directions of T4 by bar contrast, width, and speed (n = 14 cells). All responses whose mean is significantly different (see Methods) from zero are represented with filled boxes. (D_ii_) Same as D_i_ for T5 (n=17 cells). (E_i_) Baseline subtracted T4 responses to 4-LED wide PD moving bars with preferred and non-preferred contrast (same data as Ci) overlaid. Light green trace shows preferred contrast response temporally shifted to align with trailing luminance-increment edge (bottom shows schematics of bright and dark stimulus luminance levels). (E_ii_) Same as E_i_ for T5.

T4 (T5) cells are typically referred to as ON (OFF) cells as a shorthand to indicate they are strongly selective for luminance increments (decrements). Since a moving dark bar has both a leading luminance-decrement edge and a trailing luminance-increment edge, the recorded T4 responses to a dark moving bar could simply be a delayed response to the trailing edge (Fig. 1E_i_). We compared T4 responses to PD motion of bright and dark bars and noted 3 key differences:

1. in response to bright bar motion, T4 depolarization preceded hyperpolarization; in response to dark bar motion, T4 depolarization followed an initial hyperpolarization (Fig. 1 C_i_, arrowhead),
2. decay of dark bar responses were slower than for the bright bar (compare slope of green and black traces, Fig. 1E_i_), and (3) the dark bar response peak was delayed compared to bright bar peak, even after temporally aligning to the appearance of the light-increment edge (compare light green trace to black trace, Fig. 1E_i_). The same 3 differences in the response dynamics were seen when we compared responses to PD motion of dark (PC) and bright (NC) bars in T5 cells (Fig. 1E_ii_). Because of these differences, together with the selectivity for wider bars, the directionally selective responses to NC stimuli cannot be explained simply as a PC response to the trailing edge of a moving bar. The distinct characteristics of NC responses suggest contributions of additional mechanisms.

### T4 and T5 neurons have a similar structure to their non-preferred contrast receptive fields

To uncover the mechanism for generating directionally selective responses to NC stimuli in T4 and T5 cells, we mapped the ‘static’ receptive fields of both cells. Since our moving bar stimuli are composed of discrete steps (determined by the display’s LED size), we can decompose them into bar flashes presented at each position along the movement trajectory for the duration of a single step (Fig. 1A). We presented bar flashes to T4 and T5 cells with long inter-stimulus intervals randomized for contrast (bright and dark), position along the PD-ND axis, width, and duration, such that they provided no motion information.

The PC receptive field, which maps responses to bright stimuli in T4 and dark stimuli in T5, has the same overall structure previously reported [30] – depolarizing responses on the leading side of the receptive field growing towards the center, while responses on the trailing side of the receptive field display rapid depolarization followed by sustained hyperpolarization (Fig. 2A). The NC receptive field structure is distinct from the PC receptive field yet remarkably similar between T4 and T5 (Fig. 2A). For both T4 and T5, we find that NC depolarizing responses are shifted towards the trailing side, while NC hyperpolarizing responses are shifted towards the receptive field center (defined by PC responses; Fig. 2A,B). Although this receptive field structure is most evident in responses to long duration flashes of wide bars, it is also measured (with reduced magnitude) in responses to narrower bars (Fig 2C). Since the difference between moving bar responses to NC and PC stimuli hinted at different underlying mechanisms (Fig. 1E), we highlight three features of NC flash responses that differentiate them from PC responses yet are common to both T4 and T5.

**Figure 2:**
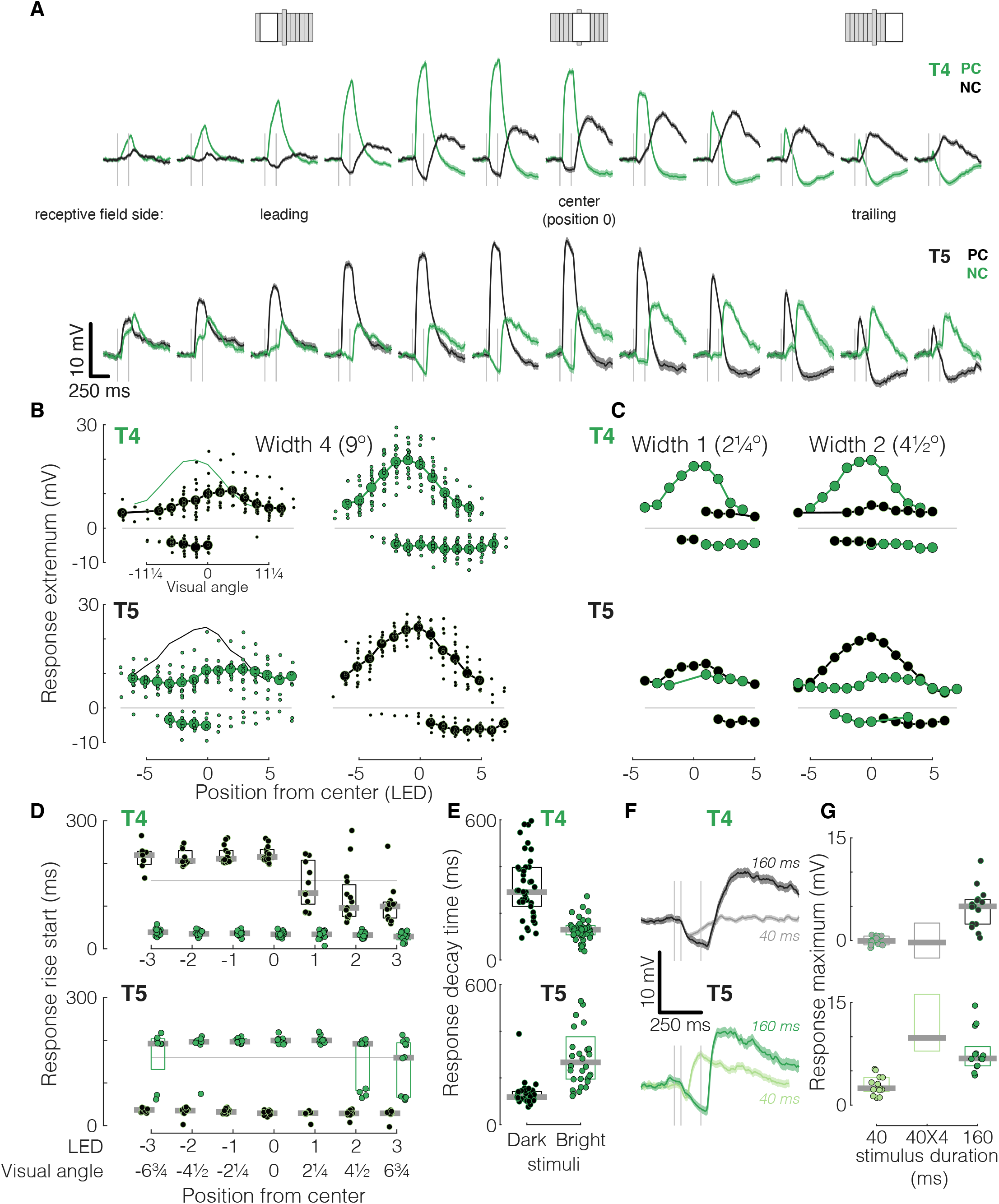
T4 and T5 neurons have a similar structure to their non-preferred contrast receptive fields. (A) Baseline subtracted responses (mean ± SEM) of T4 (middle; n=15 cells) and T5 (bottom; n=17 cells) to 160ms bright (green) and dark (black) 4-LED wide bar flashes at different position along the PD-ND axis. Gray vertical lines indicate stimulus onset and offset times. Top: Schematic depicting stimulus position within the receptive field. Subsequent panels show summary of response dynamics for this set of recordings. (B) Maximal response depolarization and hyperpolarization by position for the same stimuli as in A. Large dots represent the mean max response at each position (for positions with 4 cells or more responding), small dots the response means from individual cells. Points above (below) the zero line represent maximal depolarization (hyperpolarization). Green line in the T4 NC plot and black line in T5 NC plot represents respective PC response means for comparison. The inset axis in the top left relates the approximate visual angle along the receptive field to the 2¼° LED-defined discrete positions. (C) Same as B, only for 1- and 2-LED wide bar flash responses. Only the mean maximal responses at each position are presented. (D) Rise start time (time to reach 10% of maximum depolarization) by position for responses presented in B for T4 (top) and T5 (bottom). The stimulus appears at t = 0, and the gray horizontal lines indicate the time of stimulus offset (t = 160ms). Dots represent individual cell means, gray bars represent population median, and box edges represent population lower and upper quartiles. (E) Boxplot for the response decay times (reduction from 80% to 20% of the maximal response after maximum depolarization) for trailing side responses presented in B (positions [0:2]) for T4 (top) and T5 (bottom). (F) Mean baseline subtracted responses to non-preferred contrast 4-LED wide bar flashes presented for 40 and 160ms at a single position (−1). (G) Mean response amplitude for the same NC stimuli as in F, compared to a linear approximation (averaged over positions [−2:0]).

The first feature is prominent in the receptive field center NC responses that exhibited a strong hyperpolarization aligned to stimulus onset (Fig. 2A, first vertical line in each panel) and a depolarization following stimulus offset (second vertical line). The offset of a dark flash is accompanied by a luminance increment which is expected to evoke a depolarizing response in T4s; likewise, the bright flash offset is a luminance-decrement, the preferred stimulus for T5s. However, we find that the onset of these stimuli (signaling a luminance change opposite to the preferred one for each cell type) also evoked substantial responses. In response to NC bar onset, T4 and T5 were hyperpolarized in the receptive field center but depolarized on the trailing side. The depolarization response rise times (Fig. 2D) for PC stimuli in both T4 and T5 are position-independent and follow stimulus onset (t = 0). Conversely, rise start times for NC stimuli responses show a clear positional dependence: following stimulus offset on the receptive field leading side and center, yet following stimulus onset on the trailing side (albeit slower than for bright flashes, Fig. 2D). These results suggest specific interaction between ON and OFF signals, with T4 receiving both depolarizing and hyperpolarizing inputs induced by luminance decrements, and T5 receiving both depolarizing and hyperpolarizing inputs induced by luminance increments.

The second common feature, hyperpolarization in response to NC flashes decayed faster than hyperpolarization in response to PC flashes, is summarized for positions 0:2 (Fig. 2E). PC responses can be modeled by a combination of a fast excitatory conductance and a slow inhibitory one [29,30]. Consequently, the response decay time is likely dominated by the persistent inhibitory conductance pulling the membrane potential down. Since the hyperpolarization present in the NC flash at the receptive field center preceded the depolarizing component, we deduce that this hyperpolarization wanes faster than the slower PC hyperpolarization. These results suggest there are (at least) 2 separate sources of inhibition affecting the receptive field structure, one in response to the onset of PC stimuli and one in response to the onset of NC stimuli.

The third common feature of the NC responses is stimulus-duration-dependent offset depolarization. In response to a short dark flash, T4 cells hyperpolarized during stimulus presentation and then returned to baseline. When the same bar was presented for longer, its disappearance evoked a strong depolarization (Fig. 2F, top). The dependence on stimulus duration is not linear (Fig. 2G, paired sample t-test p < 0.001), and does not appear to rely on an intrinsic mechanism (Fig. S1). In T5 cells a similar, but smaller, stimulus-duration dependence is seen at the offset of a bright flash (Fig. 2F, bottom), exhibiting a weak sublinearity (Fig. 2G, paired sample t-test p < 0.01). Both the offset responses and the input history dependence were not found in our previous measurements of PC responses and were therefore not incorporated into our previous models [29,30].

### A unified model architecture explains the generation of both preferred and non-preferred contrast directionally selective responses in T4 and T5

Our previous work showed that a model based solely on the PC flash responses of T4 and T5 cells can predict (and thus explain) their responses to PC moving bars and moving grating stimuli [29,30]. Here, we evaluate whether a model of the PC-NC composite receptive field can explain the generation of directionally selective responses to both dark and bright moving stimuli. We established a model that allowed us to capture responses to both the onset and offset of luminance increments and decrements. We expanded our previous modelling framework with three major modifications: (1) inputs that represent stimulus offset, (2) preferred and non-preferred inputs, and (3) two additional conductances corresponding to NC inputs (Fig. 3A). We designed the model to simulate either T4 or T5 neurons, but use the T4 response to a dark bar flash (depicted in Fig. 3A, right side), to provide intuition for each of these components.

**Figure 3:**
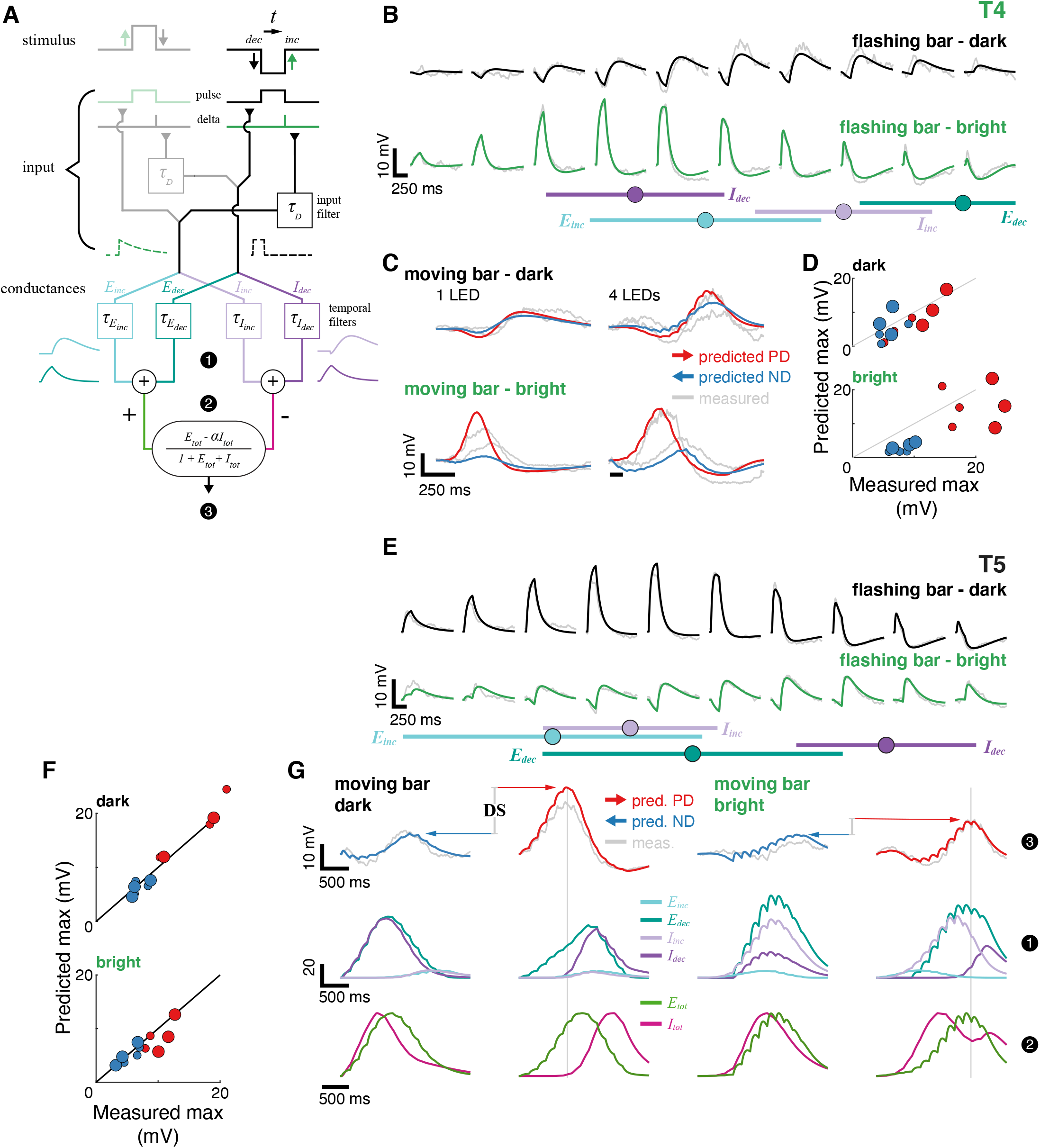
A unified model architecture explains the generation of both preferred and non-preferred contrast directionally selective responses in T4 and T5. (A) Schematic of four-conductance model used to simulate the T4 and T5 receptive field and predict responses to other stimuli. The schematic highlights the temporal aspects of the model and illustrates an example of a T4 responding to a dark (NC) stimulus. At the input stage, the stimulus generates a *pulse* signal for the duration of the light decrement and a *delta* signal marking the time of light increment (see text and Methods for details). The model includes an E-I conductance pair responding to light increments and an E-I pair responding to light decrements. Colored temporal filters represent input and conductance time-series for this specific input. The excitatory and inhibitory conductances are summed before being integrated using the biophysical nonlinearity that generates membrane voltage. Numbers match signals at each stage of the model that are shown for the T5 example in G. (B) Mean measured responses to 4-LED wide flashes of dark and bright bars at 10 different positions (in gray) compared to predicted model responses (in green and black); model was optimized to fit the averaged T4 responses (n=5 cells with cardinally aligned PD-ND axis). Bottom: gaussian spatial filters for the 4 conductances in the model (μ ± σ). (C) Mean measured responses to 1(left) and 4 (right) LED wide dark (top) and bright (bottom) bars moving in the preferred and non-preferred directions overlaid with predicted responses from the same model cell with the same model parameters as in B. (D) Peak measured responses compared to peak predicted responses for moving dark (top) and bright (bottom) bar stimuli (same widths and speeds as Fig. 1D). Marker size denotes the 2 different speeds, with larger markers representing slower movement. Marker color denotes movement direction, as in C. (E) Comparing the measured and the predicted responses, same as B but for the averaged T5 cell (n=5 cells cardinally aligned PD-ND axis). The identical model architecture was used as for T4, but since it was optimized to fit T5 responses, light decrement conductances received stronger weights. Note that the T5-optimized model has E_inc_ fed by a spatial filter on the leading side of the receptive field, to generate the leading-side bright stimulus onset component. (F) Same as D for the same averaged T5 cell as in E, presenting both dark (top) and bright (bottom) moving bar responses. (G) Example T5 moving bar responses to explain the generation of directional selectivity. The numbers on the right correspond to numbered stages in A. Top row: Mean measured responses to 4-LED wide dark (left columns) and bright (right columns) bars moving in the preferred and non-preferred directions overlaid with predicted responses from the same model cell with the same parameters as in E. Middle row: conductance traces that generated the voltage responses above. Note that the E-I increment pair has very little contribution to dark moving bar responses. Bottom row: summed total excitatory and inhibitory conductances for the 4 conditions. Gray vertical lines added to facilitate comparisons between peak voltage (top row) and the corresponding conductances.

To enable the model to respond differentially to stimulus onset and offset, we represent stimulus flashes with an input *pulse* for the duration of stimulus presentation. To account for the instantaneous luminance change that occurs at stimulus offset, we included a new *delta* input that marks stimulus offset. The duration of the preceding flash scales the amplitude of this impulse (delta function) to account for the history-dependence of the NC offset response (Fig. 2F,G), which is filtered (τ_D_ parameter, see Methods) before affecting the downstream conductances. The example dark bar flash (Fig. 3A) is composed of a luminance decrement, represented by a pulse, followed by a luminance increment, driving a delta input.

To implement both dark and bright stimuli, we split the input, such that the bright and dark input streams determine which conductance pair (increment/decrement) will receive which input type (*pulse/delta*). In our formulation the cell cannot receive both a dark and a bright input from the same position at the same time. Therefore, we can simply integrate the two parallel input streams and their resulting conductances across time (and across the spatial receptive field). The left side in the schematic (Fig. 3A, faded), conveys bright inputs and is inactive for a dark bar stimulus (see also Fig. S2). We have deliberatively not modelled complete ON and OFF pathways, both to minimize the overall number of parameters and because the details of these pathways are still under-constrained by available data.

The expanded model requires four conductances: an excitatory-inhibitory (E-I) pair responding to increments (E_inc_ and I_inc_), and an E-I pair responding to decrements (E_dec_ and I_dec_). Each pair receives the same type of input, depending on which stimulus was presented at that position. For the dark bar example, the decrement-responding E-I pair receives the *pulse* input, and the increment-responding pair receives the *delta* input; for a bright bar the input types are reversed (detailed in Fig. S2). Each conductance has its own temporal filter and a spatial filter, which is not shown in schematic (Fig. 3A and Methods).

We fit and simulated responses from a full model as well as a model where we minimized the total number of parameters by constraining some to be common or fixed (see Methods). We present results from the reduced version here (detailed results from both models in Fig. S3, S4). We optimized model parameters for T4 and T5 separately by using an iterative least-squares procedure that minimized the difference between our simulation results and the measured dark and bright bar flash responses. Since the model was fit using only responses to non-moving stimuli, later predictions of responses to moving stimuli are based solely on this ‘static’ receptive field. Notably, we use an identical model architecture for T4 and T5, and the optimization procedure achieves the appropriate luminance preference by adjusting parameters for each neuron type so that simulated responses match the corresponding measured flash responses.

As expected, the T4-optimized model’s responses were well-matched to the measured responses to both bright and dark flashes at all positions along the receptive field (Fig. 3B). Reassuringly, despite having two additional conductances, the optimization procedure converged on spatial filters for E_inc_ and I_inc_ that were similar to our prior (PC-only) T4 model: E_inc_ in the receptive field center and I_inc_ towards the trailing side (depicted by horizontal lines below traces, Fig. 3B). In the T4-optimized model, E_dec_ appears to play a minor role, fit to a spatial filter far on the trailing side (Fig. 3B) and with minimal amplitude. The I_dec_ conductance is toward the leading side and contributes the hyperpolarizing current in response to a dark bar’s onset. We next simulated moving bars responses and compared them to our recordings (Fig. 3C,D, S4). The model captures both the dynamics and the magnitude of the moving bar responses for both dark and bright bars, and, more importantly, generates directionally selective responses to both (Fig. 3G, demarcated for T5).

The T5-optimized model was fit using the identical architecture and accurately reproduces T5 responses to both bright and dark flashing bars at all positions (Fig. 3E). Since T5 prefer luminance decrements, this model has E_dec_ in the receptive field center and I_dec_ towards the trailing side (Fig. 3E, bottom). This asymmetric configuration generates a directional preference for moving dark bars [30]. The spatial filter for I_inc_ is towards the leading side and generates the hyperpolarizing current in response to the onset of a bright bar. E_inc_ is further towards the leading side and captures the weak depolarization in leading positions’ responses to bright flashing bar onset (Fig. 2A, leftmost positions). This T5-optimized model accurately predicted the magnitude of the directionally selective responses to both bright and dark moving bars (Fig. 3F, G – top row, Fig. S4). Both the T4- and T5-optimized models predict directionally selective responses to moving stimuli of both the preferred and the non-preferred contrasts.

How do the four conductances contribute to the generation of directional selectivity? We use T5 responses to slow moving bars to illustrate the mechanism. The model responses (Fig. 3G, top row) closely follow the recorded data for dark (PC) and bright (NC) bars moving in the preferred (PD) and non-preferred (ND) directions. For dark moving bars, the simulated response is dominated by two conductances: E_dec_ and I_dec_ (Fig. 3G, middle row, 2 left columns). For bright moving bars (Fig. 3G, middle row, 2 right columns), all four conductances contribute, with I_inc_ playing a much more prominent role. Considering the net effect during both dark (Fig. 3G, bottom row 1^st^ column) and bright (3^rd^ column) ND bar motion, the total excitatory and inhibitory conductances largely overlap, reducing changes in membrane potential. In response to both dark (bottom row, 2^nd^ column) and bright (4^th^ column) PD bar motion, the excitation and inhibition maxima are well-separated in time, reducing suppression of membrane depolarization. For dark stimuli, the excitation peak precedes inhibition, while for bright stimuli the excitatory conductance peaks in between the I_inc_ onset peak and the I_dec_ offset peak (similar to results of the T4-optimized model, Fig. S3, S4). For dark and bright PD bar motion, the peak membrane depolarization results from the temporal separation between the inhibitory and the excitatory conductances (Fig. 3G, vertical gray lines). This simple mechanism makes clear predictions about the range of stimulus speeds for which NC directionally selective responses can be generated. Since the excitatory and inhibitory inputs that structure the T4 and T5 receptive fields are understood to arise from physical circuit elements with fixed positions relative to each other [9], the logic of E-I overlap in ND versus E-I separation in PD can only function within a specific stimulus speed range. If a bar moves too fast, then E-I conductances will increase their temporal overlap, if it moves too slow, ND responses will be more separated (Fig. S5). Taken together, these results demonstrate that a four-conductance model optimized to reproduce the static receptive field can quantitatively account for PC and NC directional selectivity. To critically test the broader utility of our expanded model, we chose stimuli that include novel combinations of bright and dark components.

### Four-conductance model explains illusory motion percept

The mechanism we describe for generating directionally selective responses to NC moving bars should require a trailing stimulus edge, since it provides the offset signal that drives the dominant (PC) conductance pair. For the T5 example, E_dec_ and I_dec_ are the major conductances responding to both PC and NC bar motion (Fig. 3G), but for NC bar motion, they respond to the trailing edge delta input (capturing offset responses in Fig. 2A). To test this requirement, we recorded and simulated T5 responses to dark and bright moving edges (see inset Fig. 4A; no trailing luminance change, but a large return-to-baseline luminance change, lacking any directional information, occurs when edge disappears). We again find that the model, simulated with the same parameters as in Fig. 3, predicts the magnitude, dynamics and directional selectivity of the measured T5 responses to moving dark edges (Fig. 4A, left). Both the recordings and the model predictions show similar responses to a bright moving edge, featuring a prominent offset depolarization when the edge disappears (Fig. 4A, right). Importantly, these offset responses do not differentiate between the two directions. These results confirm our hypothesis that directional selectivity to NC stimuli requires a trailing luminance-change boundary, and show that moving PC, but not NC, edges drive directionally selective responses.

**Figure 4:**
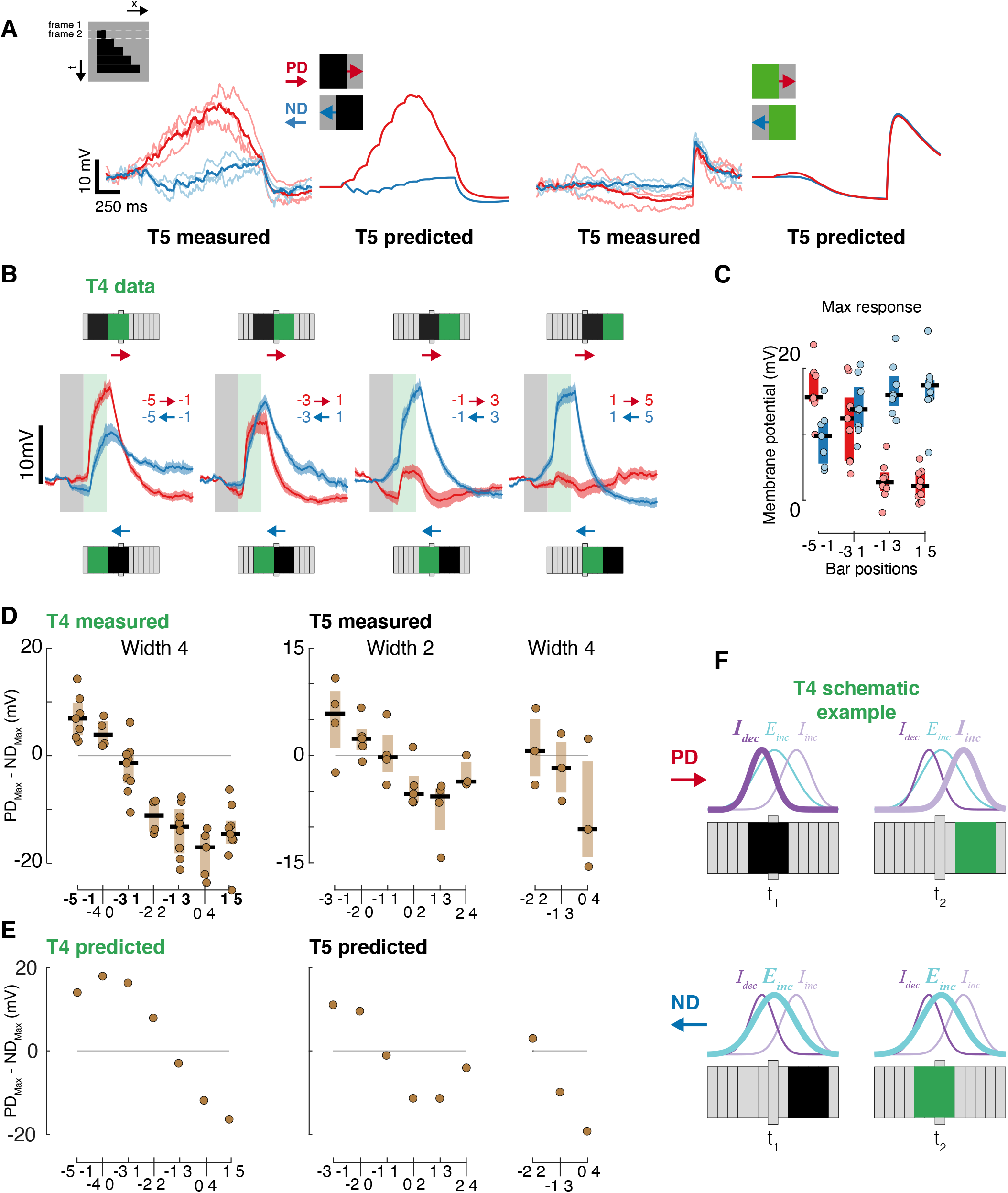
Four-conductance model explains illusory motion percept. (A) Mean baseline subtracted measured responses (n=3 individual cells in lighter shades) of T5 cells to dark and bright edges moving in the preferred and non-preferred directions, together with predicted moving edge responses from the same averaged T5-optimized model (with same parameters) as in Fig. 3. Inset shows the space-time plot for a local moving edge stimulus (moving at 160ms per LED step and for the same span as our moving bar stimuli). (B) Baseline subtracted T4 responses (mean±SEM) to dark-bright bar pairs, a minimal reverse-phi stimulus, presented sequentially along the PD-ND axis in both directions (n=7,9,8,9 cells for panels from left to right). Schematics above (below) traces depict positions for PD (ND) stimuli with respect to the center of the receptive field (extended central column). Faint gray and green rectangles demarcate presentation time of flashed bars, with each bar presented for 160 ms. (C) Boxplot summary of measured response maxima for the time series in B. Note inversion in directional preference toward the trailing side of the receptive field. (D) Boxplot summary of measured response maxima differences between PD and ND minimal reverse-phi bar-pair flashes, for T4 and T5 cells (dots represent mean responses from individual cells). The bolded labels correspond to the T4 responses for the position pairs presented in B and C. (E) Predicted response maxima differences between PD and ND bar-pair flashes for the T4 and T5 models (with same parameters) as in Fig. 3. (F) Schematic showing how the conductance model can explain the inversion in directional preference in response to a minimal reverse-phi stimulus. The spatial filters for each indicated conductance are represented above the stimulus sketches. Bold lines represent the primary conductance activated by the corresponding stimulus.

Can this model predict T4 and T5 responses to the specific ON-OFF combination that give rise to the reverse-phi illusion? In standard ‘phi’ or apparent motion stimuli, discrete sequential edge displacements are perceived as smooth motion in the direction of displacement (explains why humans enjoy movies played at only 24 frames/second). In the reverse-phi motion illusion, which has been documented in animals as diverse as flies [12], fish [15], and humans [11,33], displacement is combined with contrast inversion (dark turns bright and vice versa) resulting in motion perceived in the direction opposite to the displacement. T4 and T5 neurons have recently been shown to exhibit inverted directional responses to reverse-phi [28]. Based on our detailed characterization of T4 and T5 responses to bright and dark flashes, we asked whether the directionally inverted response to reverse-phi could also be predicted by the receptive field structure captured in our model.

A minimal version of reverse-phi illusory motion can be evoked by the sequential presentation of two adjacent bars with opposite contrasts [12,28,34]. We presented an NC bar (dark for T4, bright for T5) followed by an adjacent PC bar in multiple positions along each cell’s receptive field. Since the bar flashes appear sequentially, they signal directional information, and were presented in both directions. On the leading side of T4’s receptive field, the bar pair evoked responses consistent with the cell’s typical directional preference, despite the presence of the dark bar (Fig. 4B). However, on the trailing side, the dark-bright bar pair evoked responses with an inverted directional preference (Fig. 4B, C: [−1,3] and [1,5] pairs). We measured T5 responses to bar pairs composed of a bright (NC) bar followed by a dark (PC) one and found a similar structure. Responses to leading-side stimuli had the same directional preference as to moving PC bars, but responses showed an inverted directional preference for pairs that included trailing-side positions (Fig. 4D). This directional preference inversion was largest for wider bars presented for longer durations (Fig. 4D, S6). We used the T4- and T5-optimized models (same parameters as in Fig. 3) to simulate responses to these NC-PC bar pairs and found they accurately predicted the inversion in directional preference in both cells, including the specific relationship between the inversion and receptive field positions (Fig. 4E).

Which aspects of the receptive field structure captured by our model account for this striking inversion at the heart of the reverse-phi illusion? An NC bar (dark for example T4, Fig. 4F) in the receptive field center evokes a strong inhibitory response (I_dec_). When followed by a PC bar (bright for T4) on the trailing side, another strong inhibitory response is evoked (I_inc_). So while the bars “move” in the preferred direction, the response is strongly inhibited (Fig. 4F, top, compare to Fig. 4B PD response −1 → 3). Conversely, when the dark bar first appears on the trailing side it evokes an excitatory response (E_inc_) activated at stimulus offset. A subsequent bright bar in the receptive field center evokes a strong depolarizing response (same E_inc_ conductance) due to the stimulus onset. Therefore, while the bars “move” in the non-preferred direction, the net response is strongly excitatory (Fig. 4F, bottom, compare to Fig. 4B ND response −1 ← 3). The parsimonious model based on the high-resolution characterization of the static receptive field predicts not only PC and NC moving bar responses (Fig. 3), but also responses to moving edges (Fig. 4A), and the illusory response to PC-NC mixing reverse-phi stimuli (Fig. 4D).

### Behaviorally measured perception of reverse-phi stimuli corroborates model predictions

Since T4 and T5 cells are the major – if not exclusive – source of motion information in the fly’s visual system [35], we hypothesized that reverse-phi stimuli inducing stronger directional preference reversal in T4/T5 should also evoke stronger reversal in the fly’s behavioral response. Tethered flying flies turn in the direction of a coherently rotating grating pattern, a reaction known as the syn-directional optomotor response [36]. However, in the reverse-phi stimulus, a contrast inversion appears at every motion step (Fig. 5A, inset), to which flies respond by turning against the pattern’s rotation [12]. The receptive field mapping (Fig. 2B) and the minimal reverse-phi responses (Fig. 4, S6) showed that longer durations flashes and wider bars evoked stronger responses to NC stimuli. The tuning properties of T4 and T5 neurons should constrain the perception of dynamic stimuli integrated over larger eye regions. To test this conjecture, we presented tethered flies with gratings, comprised of bars with one of three widths, rotating at one of three speeds. We presented both standard and reverse-phi versions of these patterns in both directions. The spatial wavelength and rotation speed of the pattern are known to affect the optomotor response [37,38]. Accordingly, we find that for the standard patterns rotating at 30°/s, wider bars led to increased turning (Fig. 5B, black). Flying flies exhibited a robust reverse-phi illusion, by turning in the direction opposite to the standard motion response, with a similar effect of bar width: wider bars resulted in larger turns (Fig. 5B, red). The narrowest bar gratings presented evoked weak turns to standard motion and almost no responses to reverse-phi motion (Fig. 5C, left). Suggesting that the rotating narrow-bar standard grating was perceived as a modest level of motion, but the contrast inversions in the reverse-phi pattern were too weak to invert the motion percept. Flies responded to the wider-bar, slower standard motion patterns with reduced turning, but with increased opposite-direction turning to the corresponding reverse-phi stimuli (Fig. 5C, right). This behavioral characterization of reverse-phi responses corroborate our predictions from the electrophysiological measurements captured by our model: wider, slower bars drive stronger responses from NC conductances (T4: E_dec_ and I_dec_, T5: E_inc_ and I_inc_), which are necessary for inverting the T4 and T5 directional preference. As a consequence, slower, wider-bar gratings evoke a strong illusory percept of motion in the opposite direction, and therefore a stronger inversion of behavioral turning.

**Figure 5:**
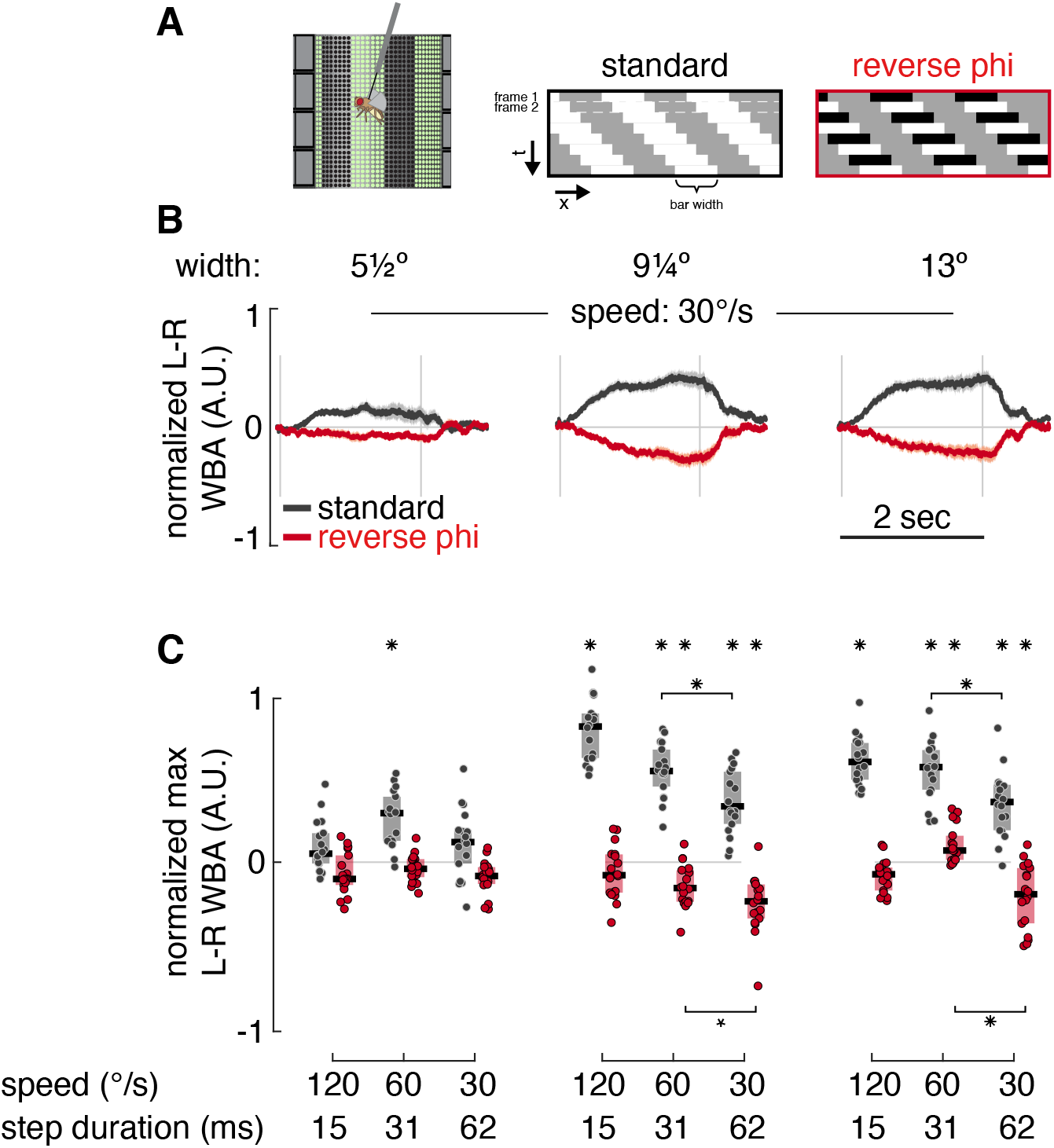
Behaviorally measured perception of reverse-phi stimuli corroborates model predictions. (A) Schematic for tethered flight arena and space-time plots illustrating the standard apparent motion and reverse-phi motion stimuli. (B) Behavioral turning responses of tethered flying flies to standard apparent motion (black) and reverse-phi motion (red) of a stimulus rotating around the yaw axis of the flies. The stimuli were presented with 3 different bar widths moving at 30°/s, in both directions, but plotted after flipping the responses to counterclockwise direction and averaging (n=18 flies, mean±SEM). Positive values indicate turning in the same direction as the direction of the stimulus. Note, that the effective angular size in the behavioral arena is different than our electrophysiological setup, so the bar widths subtend slightly different visual angles. The vertical gray lines indicate interval of stimulus motion. (C) Boxplots of maximal behavioral turning responses to apparent and reverse-phi motion stimuli with bars of the corresponding 3 widths in A, moving at 3 speeds (asterisk above boxplot – student t-test indicates a significant difference from zero, asterisk above bracket – one-sided student t-test indicates significant difference between speeds, p value, * < 0.05, * < 0.01).

## Discussion

In this study, we used whole-cell recordings from the directionally selective ON-preferring T4 cells and the OFF-preferring T5 cells and showed they also encode the motion direction of non-preferred contrast features (Fig. 1). We mapped their composite ON-OFF receptive fields and revealed a structure that is common to both T4 and T5 (Fig. 2), suggesting that both cell types receive direct, non-canonical inputs (T4 from OFF; T5 from ON). We proposed a unified model architecture to capture this composite receptive field, and showed that this model accurately predicts directionally selective responses to moving bars of either contrast (Fig. 3). The model, with no further modifications, predicted the inversion in T4’s and T5’s preferred direction in response to minimal reverse-phi motion, and also the specific receptive field locations where this inversion was measured (Fig. 4). Finally, we measured behavioral turning of tethered flying flies to reverse-phi motion and found strong evidence that the stimuli that evoked stronger NC responses in T4/T5 (wider bars, longer durations; Fig. 2) also evoked stronger behavioral inversion (Fig. 5).

### Connecting our data-driven model to motion pathway circuitry

One important goal of systems neuroscience is to connect functional measurements and the models developed to explain them with actual neural circuit mechanisms. The *Drosophila* motion-detection circuit, with a fully reconstructed connectome, genetic access to many cell types, and multiple functional studies of these cells, is an exciting system for making these connections. Nevertheless, at present we cannot simply map specific model conductances to inputs from specific upstream neurons. While both T4 and T5 neurons receive multiple PC-encoding excitatory inputs clustered around their dendritic central region [9,22] and functional studies have shown these pre-synaptic neuron types to have different temporal and spatial filters [20,24,25,39], we were able to model the PC depolarization in the receptive field center with a single excitatory conductance. Similarly, we could model a single PC inhibitory conductance, while connectomics studies have revealed at least three potential inhibitory inputs on the trailing side of the T4 dendrite [9]. It appears that T4 and T5 responses may mask additional complexity not yet uncovered, or their upstream inputs could interact (directly or indirectly) to produce simpler downstream effects.

What are the sources of the NC-mediated hyperpolarization? The T4-optimzed model places this conductance just to the leading side of the receptive field center, accounting for dark-flash hyperpolarization. Mi9 depolarizes to dark stimuli [24,25], is glutamatergic [31], and therefore likely inhibitory, and could contribute to this conductance. However, the synapses from this columnar cell type are found at the distal-most (leading) edge of T4’s dendrite [9], where we measure minimal NC-mediated hyperpolarization (Fig. 2). Further, we find similar NC hyperpolarization in our T5 measurements, but no source of NC inhibition has been described among the major T5 inputs. These discrepancies suggests that the circuit function of Mi9 remains unresolved, and that there may be undiscovered sources of NC-mediated inhibition for T4 and T5 neurons. Reconciling these model-circuit discrepancies will require better characterization of the upstream input neurons and their influence on T4 and T5, benefitting from perturbations using specific driver lines to alter neuronal function. Additionally, given the importance of both contrast changes and absolute luminance [40], assaying these cells with more complex visual stimuli should be an essential aspect of this characterization and could uncover new functions of these inputs.

### Contributions of the unified model architecture

Prior to the discovery of ON/OFF rectification by medulla neurons [18–20], nearly all models of fly motion detection, including the famous Hassenstein-Reichardt elementary motion detector, used non-rectified inputs signaling both luminance increments and decrements [41,42]. In addition to fitting the spatial and temporal tuning of insect motion detection, these classic models could also reproduce the reverse-phi illusion (a primary reason that multiplication was used for correlating offset signals). However, since we now understand that directional selectivity is computed by parallel T4 and T5 pathways, these classic models are less relevant and we must reevaluate the inverted selectivity to reverse-phi motion. Indeed, newer models have included rectified inputs [20,24,34,43], with models designed to explain the inverted directional preference to reverse-phi motion specifically including NC inputs (either as a shift in the rectification of PC inputs [10,44], or as separate inputs [45,46]). Our model differs markedly since it is subject to the strong constraints imposed by optimizing parameters to reproduce the high temporal and spatial resolution whole-cell electrophysiological measurements of T4 and T5. Additionally, our models are only trained to reproduce the static receptive field of T4 and T5 neurons, but are used to predict responses to dynamic stimuli such as moving bars, edges, and minimal reverse-phi motion. Our model is also the first (to our knowledge) in flies to incorporate stimulus offset responses (see [47] in mice). These offset components provide an intuitive explanation for why moving NC bars cause directionally selective responses while moving edges with the same contrast do not (Fig 4A), and they play a critical role in generating the inverted directional preference to reverse-phi motion (Fig. 4F). Importantly, by including offset responses in our model we can both predict the responses to reverse-phi stimuli and provide a mechanistic explanation for the illusion in flies (Fig. 4F).

### On the utility of the reverse-phi illusion

The reverse-phi illusion continues to provoke scientific interest more than half a century after it was discovered in humans [11]. In both vertebrates and invertebrates, this motion illusion appears to depend on local computations with similar spatial and temporal tuning as for standard apparent motion [12,48]. Therefore, the illusory percept is likely derived from the motion computation itself. Once challenge is that neural correlates of the reverse-phi illusion have been found throughout different visual pathways, from visual cortical neurons in primates [49] to the optic lobe in flies [44]. A previous study that proposed a related mechanism for the reverse-phi inversion seen in visual cortex, acknowledged the limitation of using data collected from diverse neuronal population to constrain models [50]. By contrast, the well-characterized *Drosophila* motion-vision circuit is an ideal system to study this illusion and its implication for motion vision, especially since the reverse-phi-mediated inversion of directional preference is found in the very same cells responsible for directionally selective motion detection [28].

Why is the origin of directional selectivity linked to an inversion in directional preference when contrast alternates? The response inversion has been proposed to facilitate more accurate motion speed estimates [28,51] or to cancel out random correlations in noisy visual inputs [12]. Our biologically constrained model of T4 and T5 responses explains the reverse-phi inversion as a consequence of an inhibitory NC component near the receptive field center. The existence of this NC inhibitory subfield is common not only to T4 and T5 neurons (despite their distinct set of upstream inputs), but also to reverse-phi-illusion-sensitive neurons in primate visual cortex [49,50]. This raises a possibly deep question: was the requirement to perceive reverse-phi motion as inverted the primary driver for the receptive field structure, or was the receptive field structure shaped in response to other factors? In other words, if perceiving reverse-phi as inverted is not the goal, why are T4 and T5 cells (and potentially other directionally selective neurons) inhibited by NC input at the center of their receptive field? Understanding this central NC inhibition should illuminate this fundamental yet underexplored aspect of seeing motion.

## Acknowledgements

We thank Aljoscha Nern for providing the driver lines and Edward Rogers for help with fly husbandry. We thank Lisa M. Ferguson for software support in the behavioral experiments. We are also grateful to Reiser lab members for support and fruitful discussions on the manuscript. This project was supported by the Howard Hughes Medical Institute.

---

**Figure S1:**
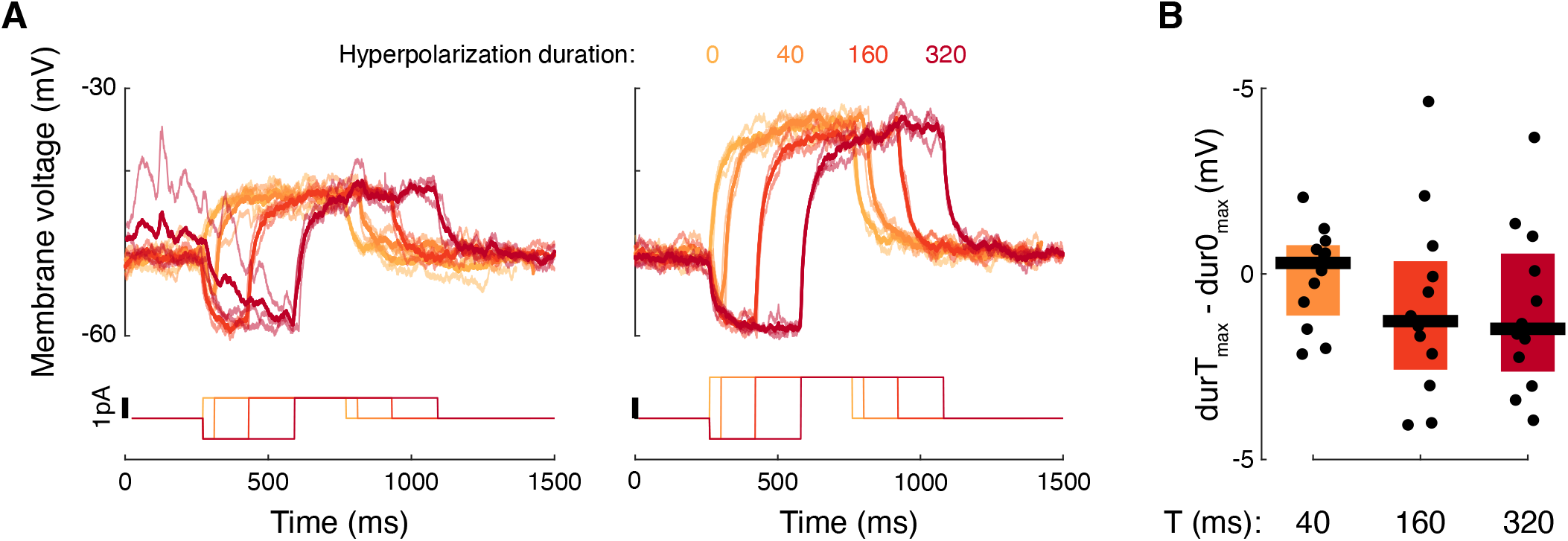
Hyperpolarizing current prior to a depolarizing pulse does not affect depolarization magnitude. Related to Figure 2. (A) Example T4 cell receiving 2 different levels of depolarizing pulses (1 and 2 pA), with preceding hyperpolarizing pulses (1 pA) of 0, 40, 160, and 320 ms durations. Bold lines are averages, thin lines are from individual repetitions of the stimulus protocol. (B) Response differences between a depolarizing current (between 0.5 to 2 pA) with preceding hyperpolarizations (between −1 to −2 pA) with the indicated duration and the same depolarizing current with no hyperpolarization (n=12 T4 cells).

**Figure S2:**
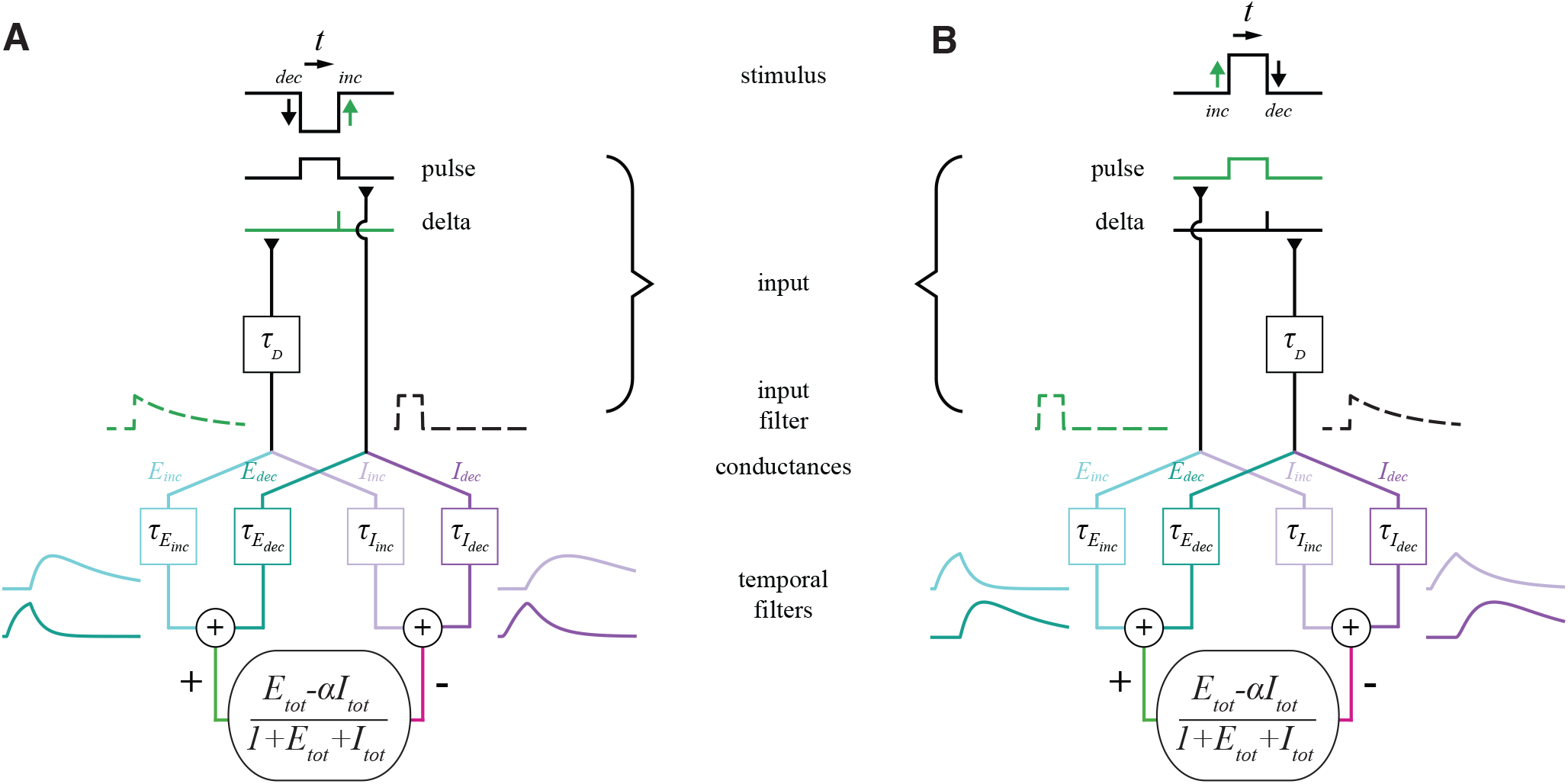
More detailed schematic for the temporal components of the T4/T5 model. Related to Figure 3. This figure shows the same as schematic as in Figure 3A, here separated into the two input conditions, by explicitly showing the inputs to a single spatial position resulting from a dark and bright bar flashes. (A) A dark flash presents a leading (in time) luminance decrement edge and a trailing luminance increment edge. The E_dec_-I_dec_ conductance pair receives the *pulse* input with the duration of the bar’s presence. The E_inc_-I_inc_ conductance pair receives a filtered *delta* input at the time of the bar’s disappearance, with a magnitude that depends on the duration of the bar’s presentation. The rest of the schematic is equivalent to the example presented in Figure 3. (B) A bright flash presents a leading (in time) luminance increment edge and a trailing luminance decrement edge. The E_inc_-I_inc_ conductance pair receives the *pulse* input with the duration of the bar’s presence. The E_dec_-I_dec_ conductance pair receives a filtered *delta* input, with the same filter as in A, at the time of the bar’s disappearance, with a magnitude that depends on the presentation duration.

**Figure S3:**
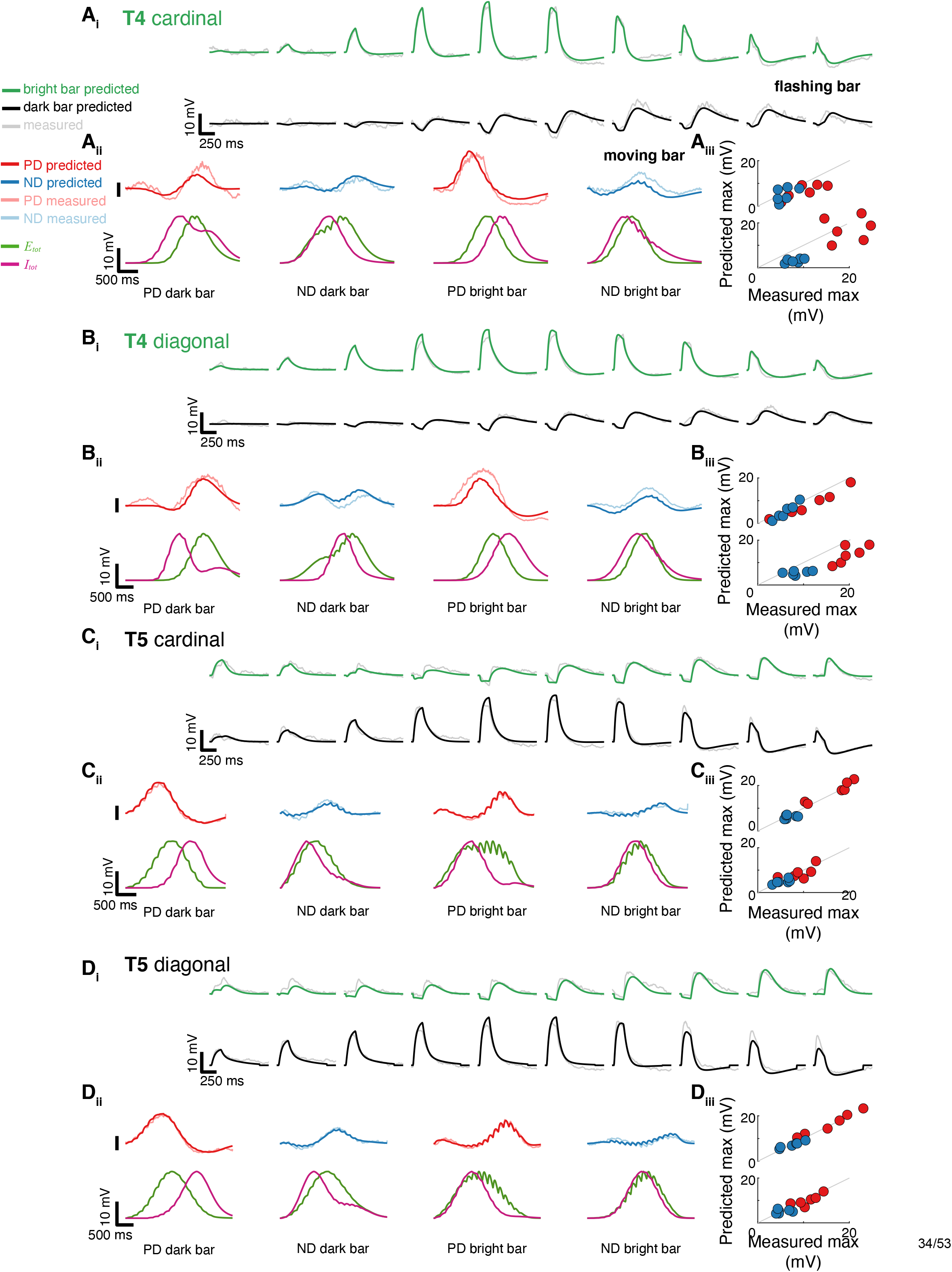
Model simulations for all averaged T4 and T5 cells from the full model. Related to Figure 3. Due to the differences in assaying the receptive fields of cells with PD-ND axis aligned to cardinal directions (on the display) compared to cells with PD-ND axis aligned diagonally (on the display), we averaged the responses for cardinal and diagonal separately, and optimized and simulated the model responses to each separately. This figure displays results from our full model, which has additional parameters (see Methods). (A_i_) Mean measured responses to 4-LED wide flashes of bright and dark bars at 10 positions (gray, same averaged response as in Figure 3, n=5 cells with PD-ND axis aligned to cardinal directions) compared to simulated model responses (green/black) for an averaged T4 cell. (A_ii_) Top: Mean measured responses to 4-LED wide dark and bright bars moving in the PD and ND at 160ms steps (14°/sec) overlaid with predicted responses from the same model cell with the same model parameters as in Ai. Bottom: summed total excitatory and inhibitory conductances for the 4 conditions above. (A_iii_) Peak measured responses compared to peak simulated responses for all moving dark (top) and bright (bottom) bar stimuli. Marker size denotes speed, with larger markers denoting slower movement. Marker color denotes movement direction, as in Aii. Same plotting conventions as in Fig. 3D. (B) Same as A for mean of T4 cells with a diagonally aligned PD-ND axis (n=9). (C) Same as A for mean of T5 cells (same averaged cell as in Figure 3, n=5) with a cardinally aligned PD-ND axis. (D) Same as A for mean of T5 cells with a diagonally aligned PD-ND axis (n=12).

**Figure S4:**
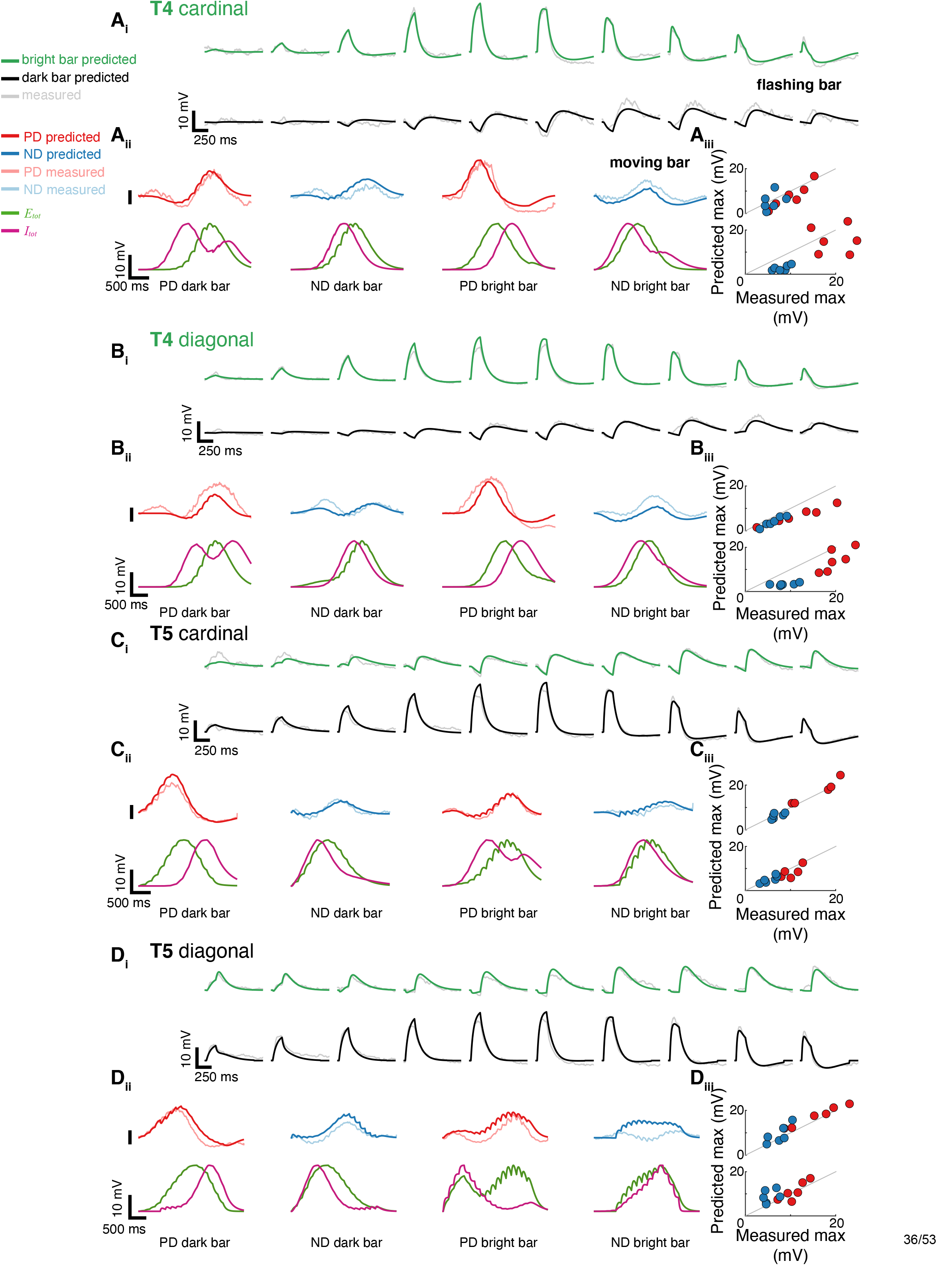
Model simulations for all averaged T4 and T5 cells from the reduced model. Related to Figure 3. Due to the differences in assaying the receptive fields of cells with PD-ND axis aligned to cardinal directions compared to cells with PD-ND axis aligned diagonally, we averaged the responses for cardinal and diagonal separately, and optimized and simulate the model responses to each separately. This figure displays results from our reduced model. Results from the reduced model optimized using T4 and T5 averages of cardinally aligned cells are presented in the main figures. (A_i_) Mean measured responses to 4-LED wide flashes of bright and dark bars at 10 positions (gray, same averaged cell as in Figure 3, n=5 cells with cardinally aligned PD-ND axis) compared to predicted model responses (green/black) for an averaged T4 cell (same parameters as in Figure 3). (A_ii_) Top: Mean measured responses to 4-LED wide dark and bright bars moving in the PD and ND overlaid with predicted responses from the same model cell with the same model parameters as in A_i_. Bottom: summed total excitatory and inhibitory conductances for the 4 conditions above. (A_iii_) Peak measured responses compared to peak predicted responses for all moving dark (top) and bright (bottom) bar stimuli. Marker size denotes speed, with larger markers denoting slower movement. Marker color denotes movement direction, as in Aii. Same plotting conventions as in Fig. 3D. (B) Same as A for mean of T4 cells with a diagonally aligned PD-ND axis (n=9). (C) Same as A for mean of T5 cells (same averaged cell as in Figure 3, n=5) with a cardinally aligned PD-ND axis. (D) Same as A for mean of T5 cells with a diagonally aligned PD-ND axis (n=12).

**Figure S5:**
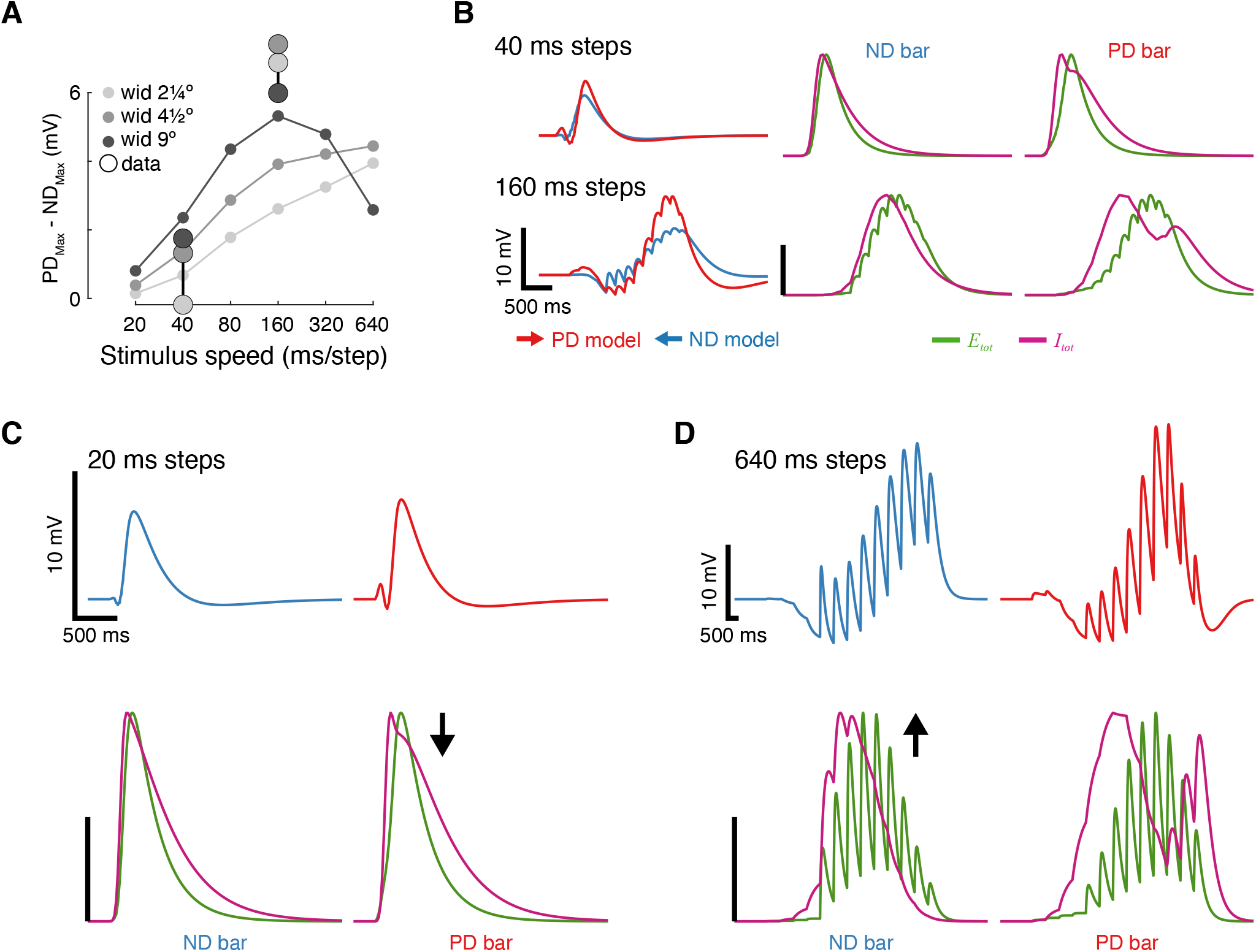
Speed tuning for non-preferred contrast moving bars. Related to Figure 3. (A) The maximum response difference for NC bars moving in the PD and ND, from the same T5-optimized model as in Figure 3. Plotted as tuning curves for model predictions for bars of the indicated widths. Large dots denote recorded data. (B) Predicted responses for 9°-wide NC bar moving at 2 different speeds. Left: predicted voltage traces for PD and ND motion. The directional selectivity of the response is larger for slower (160 ms step) moving bars. Middle: Total excitatory and total inhibitory conductances for the ND responses on the left. The membrane voltage response is reduced as a result of the temporally overlapping E and I traces. Right: Total excitatory and total inhibitory conductances for the PD responses on the left. Inhibition for the slower moving bars is decaying between the 2 peaks, transiently allowing for a more effective excitation, and thus an enhanced PD response. (C) Membrane voltage traces (top) and total conductances (bottom) for simulated PD and ND responses to a 9°-wide NC bar moving with 20 ms steps (faster than in B). Colors are as in B. Arrow near PD conductance traces indicates a reduced PD response as a result of the near fusion of the 2 inhibitory peaks (compare to B). (D) Membrane voltage traces (top) and total conductances (bottom) for simulated PD and ND responses to 9°-wide NC bar moving with 640 ms steps (slower than in B). Arrow near ND conductance traces indicates an enhanced ND response as a result of the decreased overlap between the excitatory and the inhibitory conductances.

**Figure S6:**
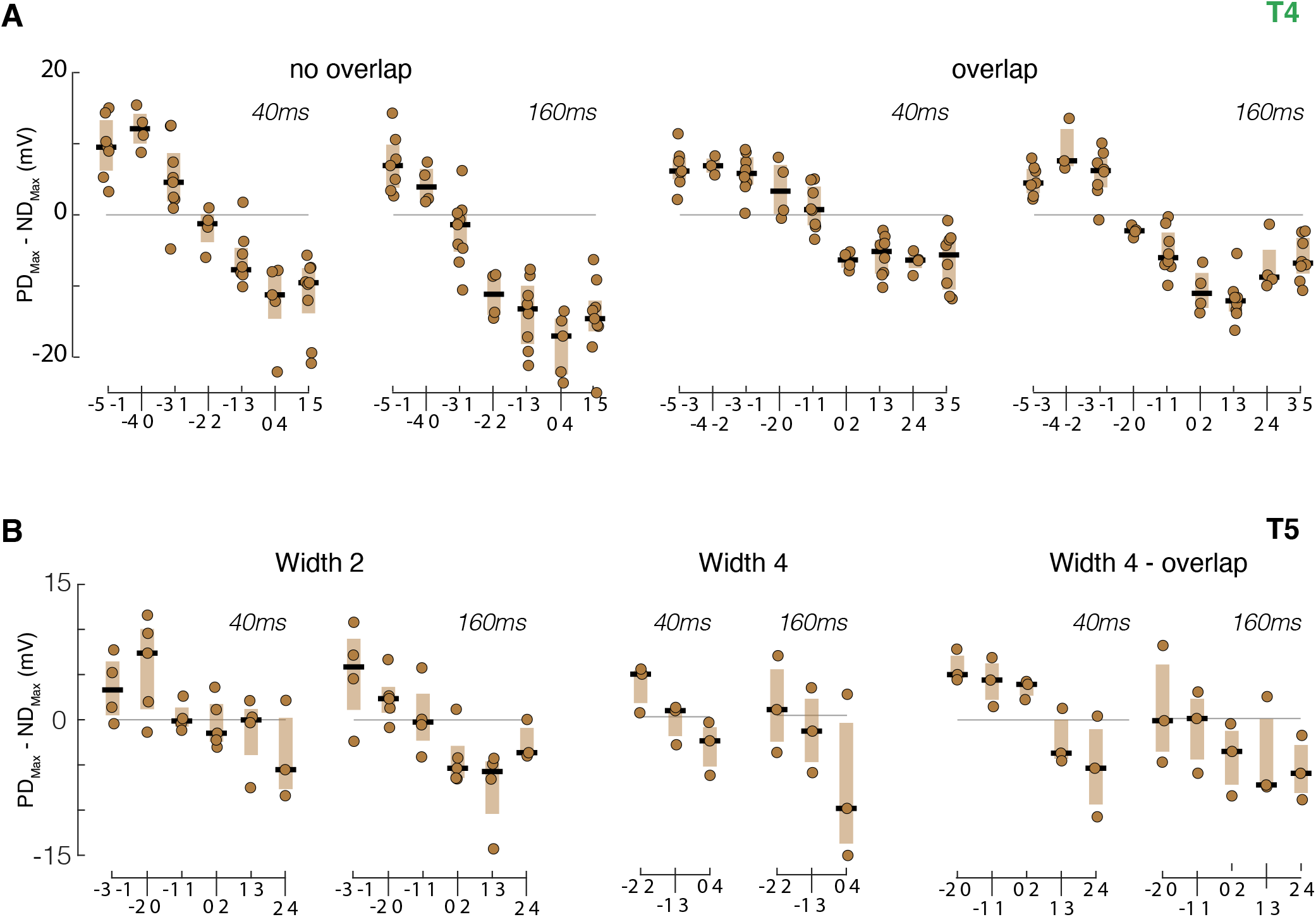
Recorded T4 and T5 responses to minimal reverse-phi stimuli depend on flash width and duration. Related to Figure 4. (A) Boxplot summary of measured response maxima differences for T4 cells presented with flashing dark-bright pairs. No-overlap stimuli with 160 ms duration are the same responses presented in Figure 4D. Overlap stimuli had the same temporal structure (1^st^ bar dark, 2^nd^ bright each presented for the duration stated above the panel) but included 50% spatial overlap between the 2 bars (also indicated by the positions, in receptive field coordinates, along the x-axis). Dots represent means from individual cells. Note that numbers for different pairs vary, since different cells were presented with different sets of pairs. (B) Boxplot summary of measured response maxima differences for T5 cells presented with flashing bright-dark pairs. No-overlap stimuli with 160 ms duration are the same response presented in Figure 4D.

## KEY RESOURCES TABLE

| REAGENT or RESOURCE | SOURCE | IDENTIFIER |
| --- | --- | --- |
| Chemicals, peptides, and recombinant proteins |  |  |
| Experimental models: Organisms/strains |  |  |
| *D. melanogaster*. T4: VT015785-p65ADZp (attP40); R42F06-ZpGdbd (attP2) | [3] | N/A |
| *D. melanogaster*. T5: VT055812-AD(attP40); R47H05-DBD(attP2) | [3] | N/A |
| *D. melanogaster*. GFP: pJFRC28-10XUAS-IVS-GFP-p10 | [2] | N/A |
| Software and algorithms |  |  |
| MATLAB 2018b | MathWorks | https://www.mathworks.com/ |
| μManager V1.4 | [6] | https://micro-manager.org/wiki/Micro-Manager |
| Other |  |  |

## STAR Methods

### RESOURCE AVAILABILITY

#### Lead contact

Further information and requests for resources and reagents should be directed to and will be fulfilled by the lead contact, Michael B. Reiser.

#### Materials availability

This study did not generate new unique reagents.

#### Data and code availability

Following the standard established in prior publications, we will make the dataset and code required to produce the major results of this study available at the time of publication. Preparing these materials is quite time-consuming, and so we will do this to correspond to the final version of the manuscript.

### EXPERIMENTAL MODEL AND SUBJECT DETAILS

Experiments were performed on 1-2 day old female *Drosophila melanogaster*. Flies were reared under 16:8 light:dark cycle at 24°C. For T5 targeted recordings we used flies with the following genotype: pJFRC28-10XUAS-IVS-GFP-p10 [2] in attP2 crossed to stable split-GAL4 SS25175 (w; VT055812-AD(attP40); R47H05-DBD(attP2)). For T4 we used pJFRC28-10XUAS-IVS-GFP-p10^44^ in attP2 crossed to stable split-GAL4 SS02344 (VT015785-p65ADZp (attP40); R42F06-ZpGdbd (attP2)). Flies of the same genotype used for targeting T4 neurons were also used for the behavioral experiments. Both genotypes were generously provided by Aljoscha Nern in Gerry Rubin’s lab [3].

### METHOD DETAILS

#### Electrophysiology

T5 recording are from cells that were included in our previous paper (n = 17 cells), and were performed as published [1], T4 recordings are from a newly acquired dataset (n = 15 cells). The experimental methods are similar to our prior manuscript [1] and will be briefly summarized below. Flies were anesthetized on ice and transferred to a chilled vacuum holder where they were mounted, with the head tilted down, to a customized platform machined from PEEK using UV-cured glue (Loctite 3972). To reduce brain motion, the proboscis was fixed to the head with a small amount of the same glue. The posterior part of the cuticle was removed using syringe needles and fine forceps. The perineural sheath was peeled using fine forceps and, if needed, further removed with a suction pipette under the microscope. To further reduce brain motion, muscle 16 [4] was removed from between the antennae.

The brain was continuously perfused with an extracellular saline containing (in mM): 103 NaCl, 3 KCl, 1.5 CaCl_2_ 2H_2_O, 4 MgCl_2_ 6H_2_O, 1 NaH_2_PO_4_ H_2_O, 26 NaHCO_3_, 5 N-Tris (hydroxymethyl) methyl-2-aminoethane-sulfonic acid, 10 Glucose, and 10 Trehalose [5]. Osmolarity was adjusted to 275 mOsm, and saline was bubbled with 95% O_2_ / 5% CO_2_ during the experiment to reach a final pH of 7.3. Pressure-polished patch-clamp electrodes were pulled for a resistance of 9.5-10.5 MΩ and filled with an intracellular saline containing (in mM): 140 KAsp, 10 HEPES, 1.1 EGTA, 0.1 CaCl_2_, 4 MgATP, 0.5 NaGTP, and 5 Glutathione [5]. 250μM Alexa 594 Hydrazide was added to the intracellular saline prior to each experiment, to reach a final osmolarity of 265 mOsm, with a pH of 7.3.

The mounted, dissected flies were positioned on a rigid platform mounted on an air table. Recordings were obtained from labeled cell bodies under visual control using a Sutter SOM microscope with a 60X water-immersion objective. To visualize the GFP labeled cells, a monochrome, IR-sensitive CCD camera (ThorLabs 1500M-GE) was mounted to the microscope, an 850 nm LED provided oblique illumination (ThorLabs M850F2), and a 460 nm LED provided GFP excitation (Sutter TLED source). Images were acquired using Micro-Manager[6], to allow for automatic contrast adjustment.

All recordings were obtained from the left side of the brain. Current clamp recordings were sampled at 20KHz and low-pass filtered at 10KHz using Axon multiClamp 700B amplifier (National Instrument PCIe-7842R LX50 Multifunction RIO board) using custom LabView (2013 v.13.0.1f2; National Instruments) and MATLAB (Mathworks, Inc.) software. Shortly after breaking in, recordings were stabilized with a small injection of a hyperpolarizing current (0-3pA) setting the membrane potential to a range between −60 to −55mV (uncorrected for liquid junction potential). Occasionally, the injected current required adjustments, but these were done prior to the acquisition of the single bar flash data. To verify recording quality, current step injections were performed at the beginning of the experiment.

#### Current injection experiments

For the experiment presented in Fig. S1 the current step injections described above were preceded with a hyperpolarizing current of different duration. This procedure was performed prior to the presentation of visual stimulation. The magnitude of the hyperpolarizing and the following depolarizing injection were adjusted manually to evoke a similar membrane voltage response between cells (~ 10 mV). Hyperpolarization current injections ranged between −1 to −2 pA, while depolarizing current injections varied between 0.5 and 2 pA.

#### Visual stimuli (electrophysiology)

The visual display was the same setup used and described in our previous paper [1]. Details are briefly summarized here. The display was constructed from an updated version of the LED panels previously described [7]. The arena covered slightly more than one half of a cylinder (216° in azimuth and ~72° in elevation) of the fly’s visual field, with the diameter of each pixel subtending an angle of (at most) 2.25° on the fly eye. Green LEDs (emission peak: 565 nm) were used, bright and dark stimuli were presented on an intermediate intensity background of ~31 cd/m^2^.

Visual stimuli were generated using custom written MATLAB code that allowed rapid generation of stimuli based on individual cell responses. In contrast to the published stimulus control system [7], we have now implemented an FPGA-based panel display controller, using the same PCIe card (National Instrument PCIe-7842R LX50 Multifunction RIO board) that also acquired the electrophysiology data. This new control system (implemented in LabView) streams pattern data directly from PC file storage, allowing for on-line stimulus generation.

To map the receptive field (RF) center of each recorded cell, three grids of flashing preferred contrast (dark for T5, bright for T4) squares were presented at increasing spatial resolution. Each flash stimulus was presented for 200ms. First, a 6 × 7 grid of non-overlapping 5 × 5 LEDs (~11°×11°) preferred contrast squares was presented (Fig. 1A). If a response was detected, a denser 3 × 3 grid with 50%-overlapping 5 × 5 LEDs (~11°×11°) preferred and non-preferred contrast squares (to verify cell polarity) was presented at the estimated position of the RF center. If a recorded cell was consistently responsive to the first two mapping stimuli, a third protocol was presented to identify the RF center. A 5 × 5 grid of 3×3 LED squares (~7°×7°) of the preferred contrast separated by 1 pixel-shifts was presented at the estimated center of the second grid stimulus. The location of the peak response to this stimulus was used as the RF center in subsequent experiments. Once the RF center was identified, a moving bar stimulus was presented in 8 directions with 80ms step duration (equivalent to ~28°/s). The bar was 9 pixels in height and 1, 2, or 4 pixels in width. When moving in the cardinal directions, the motion spanned 9 pixels. In the diagonal directions bar motion included more steps to cover the same distance (13 steps vs. 9 steps). Once the preferred direction had been estimated, bright and dark bar flashes were presented on the relevant axis for widths 1,2 and 4. To verify full coverage of RF, this stimulus was presented over an area larger than the original motion window (at least 13 positions; results in Fig. 2). Following this procedure, cells were presented with additional stimuli using the same PD-ND axis and RF center as reference frames. All stimuli were presented in a pseudorandom order within stimulus blocks. All stimuli were presented 3 times, except for single bar flashes that were repeated 5 times. The inter-stimulus interval was 500ms for moving stimuli and 800ms for single bar flashes (to minimize the effect of ongoing inhibition on the responses to subsequent stimuli).

Other presented stimuli were:

1. *Moving bar*. After identifying the PD-ND axis, bright and dark moving bar stimuli were presented along this axis using either 40ms or 160ms steps (equivalent to 56°/sec or 14°/sec respectively). Bar height was the same as for the mapping stimuli (9 LEDs) and width was either 1,2 or 4 LEDs (corresponding to 2.25°, 4.5° or 9°). Results are presented in Fig. 1, Fig. 3, Fig. S3 and Fig. S4. The moving bar stimuli presented to T4 and T5 cells were not identical. T5 cells were presented with a bar moving into and out of a presentation “window”. Meaning, a 4-LED wide bar would first appear as a 1-LED bar and would only achieve its full width once the trailing edge crossed into the stimulus window. For T4 cells, the bar’s leading edge traversed the same distance but the stimuli appeared and disappeared as full width bars.
2. *Moving edges*. Moving edge stimuli were presented in the same stimulus windows as the moving bars (and spanned the same number of steps), and with the same two values of step durations. After the edge has passed through the entire stimulus window, it disappeared, and the entire window reverted to the background levels. Results in Fig. 4A.
3. *Minimal Reverse Phi*. Bar pairs were presented such that the first bar was of the non-preferred contrast followed by a bar of the preferred contrast (bright-dark for T5; dark-bright for T4). For T5 cells, stimuli were presented in 2 different configurations. Either bars were of width 2 and the delay between the first and the second bar was adjusted to maintain a fixed speed (i.e., correcting the temporal delay to account for the spatial difference in positions), or bars were of width 4 and the second bar was presented directly after the first, regardless of positional difference. For T4 cells, only the second configuration was used since it elicited a stronger response. Results in Fig. 4 and Fig. S6. Responses presented in Fig. S6: stimuli with overlapping positions were 4-LED wide bars displayed with a 2-LED spatial overlap (bar center positions indicated along a-axis). Essentially, non-overlapping 4-LED wide bar pairs spanned 8 LEDs, while overlapping bar pairs spanned 6 LEDs, with the 2 middle LEDs inverting from non-preferred contrast to preferred contrast.

#### Behavioral experiments

Flies were reared on standard cornmeal molasses medium on a 16:8 hours light:dark cycle and were tested 0-4 hours before their subjective night to increase activity levels. All experiments were conducted on female flies from the same genotype that was used to label T4 neurons (VT015785-p65ADZp (attP40); R42F06-ZpGdbd (attP2)). Flies were cold anesthetized and tethered to a tungsten wire with UV-cured glue. Flies were given at least 30 min to recover while holding a small piece of paper, to discourage tethered flight. For these experiments the flies were placed in a center of a different visual arena spanning 270° in azimuth and ~120° in elevation. The arena consisted of 192X64 LED array with the diameter of individual LED subtending ~1.875°. The stimulus control system was the same as for the electrophysiology. The fly’s wings were illuminated from above by an IR LED (ThorLabs M850F2) and their position was monitored by an optical wingbeat analyzer (JFI Electronics Laboratory, University of Chicago, Chicago, IL, USA). Data acquisition was performed with the same control system as for electrophysiology, but with a 1KHz sampling rate.

#### Visual stimuli (behavior)

Rotating grating (standard or reverse phi) stimuli were presented in 2 second open-loop trials interleaved with 5 second “stripe fixation” closed-loop trials, during which the fly actively controlled the position of a 30° dark bar. These closed-loop trials were used to keep the fly flying and engaged in the task. Each open-loop stimulus was presented 5 times. Trials in which the fly stopped flying were excluded from the analysis. Stimuli presented in the open-loop condition were full field rotation gratings with bar widths of 5.5°, 9.25° and 13° (or 3 different spatial wavelengths with a duty cycle of 11°, 18.5° and 26° respectively), each moving with 3 angular velocities (30°/sec, 60°/sec, or 120°/sec) and 2 direction. Stimuli were presented either in the standard form, with a grating of bright over gray; or in the reverse phi form, with a bright (dark) over gray inverting to dark (bright) over gray with each motion step (see Fig. 5A).

#### T4/T5 neuron model

We modeled the membrane potential responses of T4 and T5 neurons with a single-compartment conductance-based neuron model [1,8], whose dynamics are described by

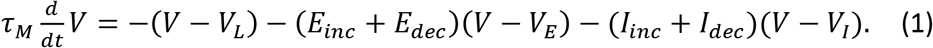

The model has four conductances: One pair of excitatory (*E_inc_*) and inhibitory (*I_inc_*) conductances respoding to luminance increments, and a second pair of conductances, *E_dec_* and *I_dec_*, for luminance decrements. All the conductances are measured in units of leak conductance. The reversal potential for the leak, excitation, and inhibition are denoted by *V_L_, V_E_*, and *V_I_* respectively.

We examined the model dynamics in the limit of small neuronal integration time *τ_M_* [1,8]. With this approximation, the dynamics of Equation 1 become

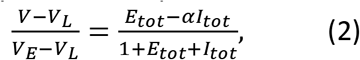

where *E_tot_* = *E_inc_* + *E_dec_* and *I_tot_* = *I_inc_* + *I_dec_* are the the total excitatory and inhibitory conductances respectively, and 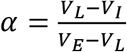.

Each of the four conductances is the sum of local contributions from receptive field locations along the PD-ND axis (*f_j_*(*t, x*),*j* = *E_inc_, I_inc_, E_dec_, I_dec_*), weighted by a spatial receptive field amplitude, described by a Gaussian. For instance, the luminance increment excitatory conductance is described by

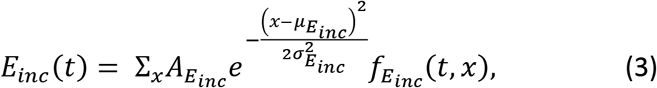

where *A_E_inc__, μ_E_inc__*, and *σ_E_inc__* denote respectively the amplitude, center, and size of the Gaussian spatial receptive field. The receptive field locations are indexed by the location *x* along the PD-ND axis (relative to the location corresponding to the empirically measured center of the receptive field of a neuron, *x* = 0). Spatial locations *x* are discretized on a uniform grid with a spacing corresponding to the smallest width of flashed bars used in the experiment. Similarly, for the remaining conductances:

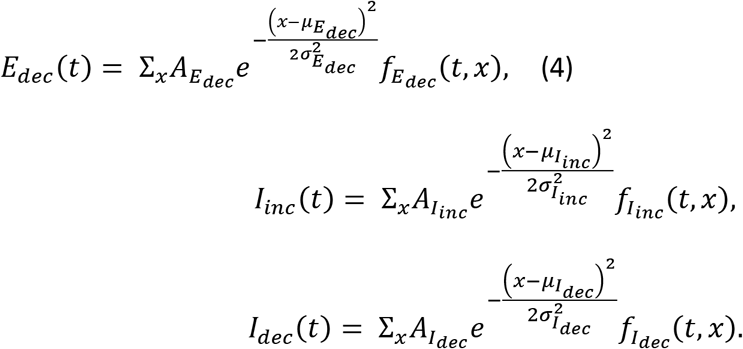

The time course of the local contribution to the conductances, *f_j_*(*t, x*), is the output of two linear temporal filters in series, with time constants *τ_j,rise_* and *τ_j,decay_*

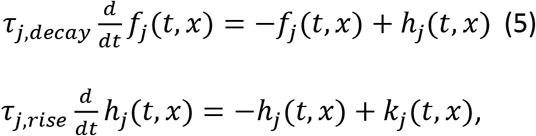

where *k_j_*(*t, x*) is a function of the visual input that depends on the stimulus type (i.e. dark or bright).

#### Input dynamics

Here we describe the model of visual input processing upstream of the T4 and T5 model neurons. For bright stimuli, the input to the temporal filters for the light increment conductances is simply

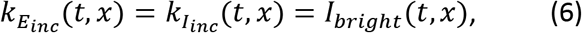

The external input *I_bright_*(*t, x*) assumes a value of 0 or 1 depending on whether a bright stimulus is absent or present at location *x* and time *t*.

For light decrement conductances, the input is a filtered delta pulse centered at the time of bright stimulus offset 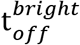:

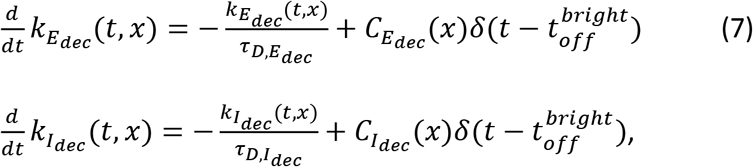

where *τ_D,E_dec__* and *τ_D,I_dec__* are the time constants of the filters. The amplitudes of the delta, *C_E_dec__*(*x*) and *C_I_dec__*(*x*), are functions of the bright stimulus duration at location *x* (*d_bright_*(*x*)). The experimental data suggests that these are nonlinear functions. In the experiments, however, we do not have a sufficiently dense sampling of possible stimulus durations to completely characterize this nonlinearity. Hence, we chose a simple rectified-linear function (ReLU):

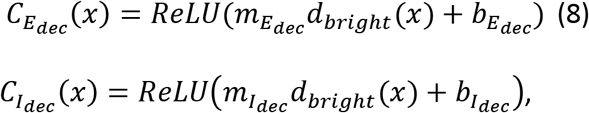

with parameters *m, b*.

For dark stimuli, light decrement conductances are directly driven by the presense of the dark stimulus *I_dark_*(*t,x*), while light increment conductances receive a filtered delta pulse at the time of the dark stimulus offset 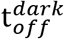:

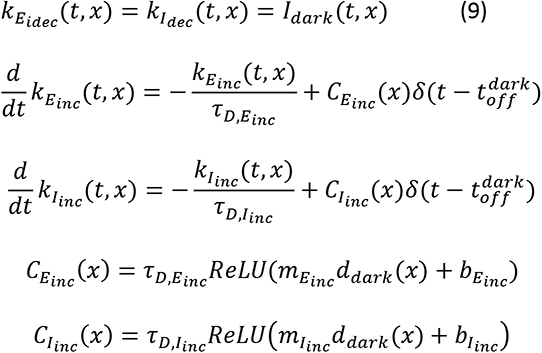

#### Model optimization

We present two version of the model: a version we have named the ‘full model’ that contains independent parameters for each conductance and was optimized using both single bar flashes and moving bars (for reasons that will be discussed below), and a version named the ‘reduced model’ in which *τ_D_* is the same for all the conductances, *τ_rise_* is fixed at 1 for the excitatory conductances, and the model was optimized using only single bar flashes (see Tables 1 and 2). Early iterations of our optimization procedure have shown these were reasonable constraints that incurred only a modest price at the loss function (Compare Figures S3 and S4). In the reduced model (Fig. 3 and Fig. S4), parameters are randomly initialized within the bounds in Table 1 and optimized to minimize the mean squared error (MSE) between simulated and measured voltage responses to PC and NC single bar flash stimuli of width [2,4] and duration [40ms, 160ms]

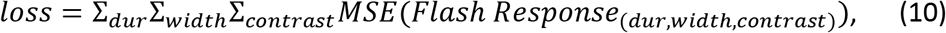

where each stimulus condition was given equal weight in the cost function.

**Table 1.** Reduced Model

| Parameters | $E_{inc}$ | $I_{inc}$ | $E_{dec}$ | $I_{dec}$ | units |
| --- | --- | --- | --- | --- | --- |
| **A** | [0,10] | [0,10] | [0,10] | [0,10] | unitless |
| $\mu$ | [-7,7] | [-7,7] | [-7,7] | [-7,7] | R.F. position |
| $\sigma$ | [.1,5] | [.1,5] | [.1,5] | [.1,5] | R.F. position |
| $\tau_{rise}$ | 1 | [1,600] | 1 | [1,600] | ms |
| $\tau_{decay}$ | [1,600] | [1,600] | $\tau_{decay,Einc}$ | [1,600] | ms |
| **M** | [0,5] | [0,5] | [0,5] | [0,5] | 1/ms |
| **b** | [-10,10] | [-10,10] | [-10,10] | [-10,10] | unitless |
| $\tau_D$ | [1,600] | $\tau_{D,Einc}$ | $\tau_{D,Einc}$ | $\tau_{D,Einc}$ | ms |

The full model (Fig. S3) parameter bounds are given in Table 2, and this model was optimized with PC and NC stimuli for both single bar flashes and moving bar stimuli of width [2,4], duration [40ms, 160ms]. Moving bars were added to the optimization procedure in an attempt to increase the density of duration sampling for equation 8. For a single position in the receptive field, bars of different widths moving at the same speed are equivalent to flashes of different durations (e.g. width 2 bar moving at 40 ms steps is equivalent to an 80 ms flash).

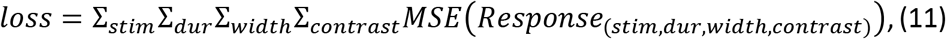

**Table 2.** Full Model

| parameters | $E_{inc}$ | $I_{inc}$ | $E_{dec}$ | $I_{dec}$ | units |
| --- | --- | --- | --- | --- | --- |
| **A** | [0,10] | [0,10] | [0,10] | [0,10] | unitless |
| $\mu$ | [-7,7] | [-7,7] | [-7,7] | [-7,7] | R.F. position |
| $\sigma$ | [.1,5] | [.1,5] | [.1,5] | [.1,5] | R.F. position |
| $\tau_{rise}$ | [1,600] | [1,600] | [1,600] | [1,600] | ms |
| $\tau_{decay}$ | [1,600] | [1,600] | [1,600] | [1,600] | ms |
| **m** | [0,5] | [0,5] | [0,5] | [0,5] | 1/ms |
| **b** | [-10,10] | [-10,10] | [-10,10] | [-10,10] | unitless |
| $\tau_D$ | [1,600] | [1,600] | [1,600] | [1,600] | ms |

All simulated responses were delayed by 30ms to match the measured transmission latencies from the omatidia to T4/T5 neurons.

The optimization for both models was performed with the MATLAB function *fminconQ* and the default interior-point algorithm.

In Figures 3 and 4 and Supplementary Figures 2, 4, and 5 we report the simulated responses from these optimization procedures. The parameters used in those simulations have been chosen among the top 1% optimization solutions based on the losses described above. Within these solutions, we selected the parameter sets resulting in the best match (smallest MSE) with moving bar stimuli not used in the loss functions for the reduced model.

### QUANTIFICATION AND STATISTICAL ANALYSIS

#### Analysis (electrophysiology)

Analysis in this paper followed similar procedure as in our previous papers [1,8] with certain necessary modification. All data analysis was performed in MATLAB using custom written code. Since the cells’ baseline was typically stable, we included only trials in which the mean pre-stimulus baseline did not differ from the overall pre-stimulus mean for that group of stimuli by more than 10 mV. We also verified that the pre-stimulus mean and overall mean for that trial did not differ by more than 15 mV (or 25 mV for slow moving bars, due to their strong responses). Responses were later aligned to the appearance of the bar stimulus and averaged (or the appearance of the bar in the central position in case of the 8-orientation moving bar).

#### Determining PD

After presenting the cell with 1- and 2-LED-wide bars moving in 8 different direction at 80 ms per LED step (speed that was optimal for determining directional selectivity for T4 cells, [8]), the preferred direction for the cell was determined by a visual estimate of the responses to determine the middle of the relatively wide range of large responding directions. The preferred direction was determined using only preferred contrast stimuli. Due to the structure of our display system (with LEDs organized in a rectangular grid), motion along diagonal directions included more steps when compared to motion along cardinal directions that span the same distance (13 rows of LEDs compared to 9). Accordingly, the responses are slightly difference, and we therefore, separated the cells into groups of either cardinally-aligned PD-ND axis, or diagonally-align PD-ND axis. When presenting average responses to moving stimuli, we present results from the diagonally-aligned T4 and T5 cells in Fig. 1, and the cardinally-aligned cells in Fig. 3. Model predictions presented in the main figures are always from T4- and T5-fits that were optimized to the cardinally-aligned cells. Figures S3 and S4 show predications from models that were optimized to each one of the four groups independently.

#### Single Position Flash Response – depolarization

responses were defined as the 0.990 quantile (a robust estimate of the max) of the response during the ‘response window’ (defined below). If the response magnitude did not exceed 2.5 standard deviations of the pre-stimulus baseline (during a 200 ms window preceding the stimulus), the response was defined as zero. For 2- and 4-LED-wide bars the threshold was 2.7 and 2.9 standard deviations, since the responses were stronger. Due to the difference in dynamics for preferred and non-preferred contrast responses, the response window was defined as a function of both stimulus duration and stimulus contrast (200 ms for PC and 375 ms for NC + flash duration). Used in Fig. 2B-E.

#### Single Position Flash Response – hyperpolarization

same as above only the response window time was defined differently. For PC stimuli, the window was 800 ms + flash duration (since the hyperpolarization appears after the depolarization). For NC stimuli, the same window as the depolarization was used, since NC stimuli induced onset hyperpolarization. In addition, lower standard deviation thresholds were used (1.5, 1.7, and 1.9), due to lower magnitude of hyperpolarization. Used in Fig. 2B-E.

#### Rise start time

Only presentations in which an average response was detected as depolarizing were used for this calculation. Start time was defined as the time from stimulus presentation (after correcting for arena delay) to 10% of the of the value of the maximal response for that position. Used in Fig. 2D

#### Decay time

Same criteria was used as for rise time calculation. Decay time was defined as the temporal difference between reaching 80% and then 20% of the maximal response (after the response peak). Used in Fig. 2E.

#### Analysis (Behavior)

All data analysis was performed in MATLAB using custom written code. Responses were quantified as left minus right wing-beat amplitude, a measure which has been shown to be proportional to yaw torque [9]. Turning responses were plotted for each individual fly and acceptable symmetry was verified between left and right turns. Next, responses to both directions of the same stimulus condition were combined into an average after inverting the sign of responses to counterclockwise moving stimuli (standard and reverse phi). For each fly responses were normalized to the 0.95 quantile of the strongest single stimulus response. Mean responses were calculated by taking the average left minus right (turning) value within the 1-2 sec response window. Used in Fig. 5

#### Statistics

To determine whether mean responses in Fig. 1D were (statistically) significantly different from zero, response groups were first tested for normality using the Lilliefors test, and then tested for a significant difference from zero using the one-sided Student’s *t*-test. To test for differences between responses to long flashes and their equivalent linear sums in Fig. 2G, a paired two-sided Student’s t-test was performed. Finally, to test for the significance of behavioral responses in Fig. 5B, a two-sided t-test was performed to determine whether responses are different from zero, and a paired one-sided t-test was performed to determine whether responses to 60°/s and 30°/s moving grating differed. No statistical methods were used to pre-determine sample sizes, however our sample sizes are similar to those reported in previous publications [10–12]. Data collection and analysis could not be performed blind to the conditions of the experiments.

#### Data Plotting Conventions

All boxplots presented were plotted with these conventions: box represents upper and lower quartile range, line represents median, whiskers were omitted, and in certain cases (that are clear from context) individual data points are overlaid on the box.

